# RBPMS2 is a conserved regulator of alternative splicing that promotes myofibrillar organization and optimal calcium handling in cardiomyocytes

**DOI:** 10.1101/2021.03.08.434502

**Authors:** Alexander A. Akerberg, Michael Trembley, Vincent Butty, Asya Schwertner, Long Zhao, Manu Beerens, Xujie Liu, Mohammed Mahamdeh, Shiaulou Yuan, Laurie Boyer, Calum MacRae, Christopher Nguyen, William T. Pu, Caroline E. Burns, C. Geoffrey Burns

**Author notes:** Correspondence: Caroline and C. Geoffrey Burns,. 300 Longwood Ave., Boston MA 02115.

## Abstract

**Rationale:** The identification of novel cardiomyocyte-intrinsic factors that support heart function will expand the number of candidate genes and therapeutic options for heart failure, a leading cause of death worldwide.

**Objective:** To identify and characterize conserved regulators of cardiomyocyte function.

**Methods and Results:** We report that the RNA-binding protein RBPMS2 is required for myofibril organization and the regulation of intracellular calcium dynamics in both zebrafish embryos and human induced pluripotent stem cell-derived cardiomyocytes (hiPSC-CMs). A differential expression screen in zebrafish uncovered enrichment of *rbpms2* paralogs, *rbpms2a* and *rbpms2b,* in the myocardium. Double knock-out (*rbpms2-*null) embryos suffer from compromised ventricular filling during the relaxation phase of the cardiac cycle, which significantly reduces cardiac output. Whole transcriptome sequencing and validation studies revealed differential alternative splicing of several genes linked to cardiomyopathies in humans, including *myosin binding protein C3* (*mybpc3*) and *phospholamban* (*pln*), consistent with a role in causing the observed ventricular deficiencies. Further, *RBPMS*2-null hiPSC-CMs exhibit myofibril and calcium handling defects that are highly analogous to those observed in the *rbpms2-*null zebrafish ventricle.

**Conclusions:** Taken together, our data identify *RBPMS2* as a conserved and essential regulator of alternative splicing that is required for myofibrillar organization and optimal calcium handling from zebrafish to humans.

## INTRODUCTION

Heart failure remains a common, costly, and growing public health burden ^1^. While lifestyle choices that cause hypertension and myocardial infarction are substantial contributors to heart failure, inherited and *de novo* genetic mutations that cause primary diseases of the heart muscle, including hypertrophic cardiomyopathy (HCM) and dilated cardiomyopathy (DCM) are also important instigators of disease ^2,3^. Causal mutations for HCM and DCM commonly reside within two classes of genes, those encoding components of the sarcomere, the basic contractile unit of cardiac muscle, and those encoding factors that regulate calcium handling, another critical determinant of cardiac contractility ^2,4–6^. While the pathogenesis of mutations affecting sarcomeres and calcium handling is well-established in HCM and DCM, a significant percentage of cases remain idiopathic, because clinically recognized cardiomyopathy susceptibility genes remain mutation free in targeted sequencing panels. Therefore, ongoing efforts to identify novel molecular activities required for optimal myocardial contractility will highlight novel candidate genes for primary cardiomyopathies.

Alternative mRNA splicing has emerged in recent years as an important regulator of myocardial contractility in humans. Disruptions in the RNA-recognition motif (RRM)-containing protein RBM20 were recently discovered to cause dilated cardiomyopathy (DCM) by regulating alternative splicing of TITIN (TTN), the largest protein component of the sarcomere ^7–9^. TTN transcripts within the heart can be separated into three distinct classes, each of which codes for a unique combination of functional domains ^10^. Loss of Rbm20 in animal models results in aberrant splicing of TTN, leading to changes in ventricular morphology and diminished contractility in a manner consistent with observations in DCM patients. Clinical studies have further confirmed that genetic variation in the Rbm20 locus is associated with a higher risk of cardiomyopathy ^11,12^. Alternatively, spliced transcripts of calcium/calmodulin-dependent kinase II delta (CAMK2D) were also shown to contribute to increased risk of arrhythmogenic cardiomyopathy in the absence of Rbm20 ^7^, demonstrating roles for Rbm20 in the regulation of myocardial calcium handling and action potential propagation. Similar to Rbm20, another RRM-containing protein, termed Rbm24, was shown to regulate the splicing of a subset of genes encoding components of the sarcomere, including TTN, within postnatal mice^13^. Initial screens for Rbm24 mutations in human cardiomyopathy patients suggest a potential link to disease, however, these mutations were found to be relatively rare in the disease population ^14^. Together these data represent the first etiological links between splicing factors and cardiomyopathy, thereby underscoring the importance for further research to identify the full array of splicing factors that regulate cardiac form and function.

The importance of alternative splicing in the heart is further evidenced by the abundance of isoform variation within cardiac-specific transcripts. Furthermore, many genes with established roles in cardiomyopathy are known to produce multiple distinct isoforms within the heart. For example, the cardiac troponin t (*TNNT2*) gene is alternatively spliced in a regulated pattern during cardiac development and aberrant splicing can lead to cardiomyopathy in animal models ^15–17^. Other cardiomyopathy genes such as tropomyosin (*TPM1*) and myosin binding protein C3 (*MYBPC3*) are predicted to be alternatively spliced in a similar fashion, however the importance of these splicing events and the identity of the factors that govern them remain unknown.

Here, we identify the RRM-containing protein Rbpms2 (RNA binding protein with multiple splicing 2) as a crucial regulator of alternative splicing within the developing myocardium. We show that Rbpms2a and Rbpms2b redundantly promote ventricular contractility within the developing zebrafish heart. Further, we show that these Rbpms2 proteins regulate the splicing of numerous genes that are associated with cardiac sarcomere assembly and ion transport, including transcripts for known disease genes *mybpc3* and *phospholamban* (*pln)* ^18,19^. We then demonstrate that *rbpms2*-null animals exhibit sarcomere disorganization and calcium handling deficits that are consistent with the misregulation of Rbpms2 targets.

## METHODS

### Data availability

All data and materials will be made publicly available upon publishing. RNA-sequencing datasets summarized in Tables S1-5 were deposited into the NCBI GEO repository under accession numbers GSE167416 and GSE167414.

### Zebrafish husbandry and strains

Zebrafish were bred and maintained following animal protocols approved by the Institutional Animal Care and Use Committees at Massachusetts General Hospital and Boston Children’s Hospital. All protocols and procedures followed the guidelines and recommendations outlined by the Guide for the Care and Use of Laboratory Animals. The following zebrafish strains were used in this study: *TgBAC(−36nkx2.5:ZsYellow)*^*fb7* 20^, *Tg(myl7:GFP)^fb1^* [formerly *Tg(cmlc2:GFP)^fb1^]* ^21^*, Tg(myl7:NLS-eGFP)^fb18^* [formerly *Tg(cmlc2:nucGFP)^fb18^]* ^22^*, Tg(myl7:actn3b-EGFP)^sd10^* [formerly *Tg(cmlc2:actinin3-EGFP)]* ^23^*, rbpms2a^chb3^* (this study), and *rbpms2b^chb4^* (this study).

### Generation and genotyping of *rbpms2a^ch3^* and rbpms2b^*ch4*^ mutant alleles

Customized transcription activator-like effector nucleases (TALENs) ^24–26^ were used to induce double-stranded breaks in regions of the *rbpms2a* and *rbpms2b* loci that encode the RNA-recognition motifs. TALENs were designed to bind the sequences 5’-TGCCTGTTGACATCAAGC-3’ (+) and 5’-TTGAAGGGTCTGAACAGC-3’ (-) in exon 2 of *rbpms2a* and 5’-TGCCAACAGATATCAAAC-3’ (+) and 5’-TTAAATGGTCGAAATAGC-3’ (-) in exon 2 of *rbpms2b*. TALEN constructs were constructed as described ^26^. Messenger RNAs encoding each TALEN pair were produced in vitro and co-injected into one-cell stage zebrafish embryos as described ^25^ Germline transmission of TALEN-induced mutations was detected using fluorescent PCR and DNA-fragment analysis as described ^27^. The nucleotides deleted in the mutant alleles *rbpms2a^chb3^* and *rbpms2b^chb4^* are shown (Fig. S4A,B). The primers utilized to distinguish wild-type *rbpms2a* and *rbpms2b* from the mutant alleles *rbpms2a^chb3^* and *rbpms2b^chb4^* by fluorescent PCR and DNA-fragment analysis are shown (Table S5). The wild-type alleles produce amplicons of 97 base pairs (bp)(*rbpms2a*) and 76bp (*rbpms2b*), whereas the mutant alleles produce amplicons of 83bp (*rbpms2a^chb3^*) and 74bp (*rbpms2b^chb4^*).

### hiPSC maintenance and genome editing

All hiPSC lines were maintained at 37°C, 5% CO_2_ in Essential 8 medium (Cat. # A1517001, Thermo Fisher Scientific) and passaged every 3 – 4 days using Versene (Cat. # 15040066, Thermo Fisher Scientific). Culture dishes were pre-coated with 0.5% GelTrex matrix solution (Cat. # A1413302, Thermo Fisher Scientific). CRISPR/Cas9 genome editing was performed to create isogenic lines. Briefly, guide RNA for *RBPMS2* (sequence 5’-GTCTTGCAGTGAGCTTGATC-3’) was synthesized using the T7 MegaShortScript transcription kit (Thermo Fisher Scientific). We previously described methods ^28^ to generate a doxycycline - inducible Cas9 line in the WTC-11 iPSC background (Coriell Institute). Targeted integration of Cas9 into the AAVS1 locus was performed using Addgene plasmids #73500 and #48139. This WTC-11 AAVS1^Cas9/+^ hiPSC line was treated with doxycycline (2μg/ml) for 16 hours prior to transfection to induce expression of an NLS-SpCas9 fusion protein. After induction, approximately 0.5 x10^6^ cells were transfected with 10μg guide RNA using an Amaxa nucleofector (Lonza Technologies). Cells were seeded at a low density to pick single clones. Positive clones were confirmed by Sanger and amplicon sequencing. Stem cell marker expression was validated in isolated cell lines by flow cytometry using anti-SSEA4-FITC (Cat. # 130-098-371, Miltenyi Biotec).

## RESULTS

### Differential expression analysis of FACS-purified zebrafish anterior lateral plate mesodermal cells identifies cardiomyocyte-restricted expression of rbpms2a and rbpms2b

Lineage-restricted genes often perform specialized and essential functions in the development, maintenance, or function of the populations they mark. To generate a candidate list of zebrafish genes with essential activities in cardiopharyngeal progenitors or embryonic cardiomyocytes, we sought to identify mRNAs with restricted expression to the subset of anterior lateral plate mesodermal (ALPM) cells marked by the *nkx2.5:ZsYellow* transgene. This population contains cardiopharyngeal progenitors for multiple cell types including second heart field-derived ventricular cardiomyocytes, three outflow tract (OFT) lineages, a subset of pharyngeal muscles, and endothelium of the pharyngeal arch arteries ^20,29–32^. It also contains early-differentiating ventricular cardiomyocytes derived from the first heart field ^29,31,32^.

To isolate ALPM cells labeled with the transgene, *Tg(nkx2.5:ZsYellow)* embryos were dissociated to single cells at a developmental stage when ZsYellow fluorescence is restricted to the ALPM [14-16 somites stage (ss); Fig. 1A]. Dissociated cells were separated by FACS into ZsYellow-positive and ZsYellow-negative subpopulations before being subjected to RNA-sequencing and differential expression analysis (Fig 1A; Fig S1A-C). From this, we identified 1848 protein-coding RNAs significantly enriched in the ZsYellow-positive population [fold change (FC)>1.5; adjusted p-value<0.05; Fig. 1A; Fig. S1D; Table S1]. Among them were several previously characterized lateral plate mesodermal markers that encode transcription factors essential for vertebrate cardiopharyngeal development including *nkx2.5*, *hand2*, *gata4/5/6*, *mef2ca/b*, *tbx1/5a/20*, *isl1*, and *fgf8a/10a/10b* (Fig. S1D; Table S1). Due to the presence of early-differentiating cardiomyocytes in the ZsYellow-positive population, we also identified numerous genes encoding sarcomere components [*cmlc1*, *myl7* (*cmlc2*), *myh7* (*vmhc*), *myh6* (*amhc*), *acta1b*, *actc1a*, *actn2b*, *tnni1b*, *tnnt2a, tnnc1a, tpm4a, ttn.2*], calcium handling proteins [*slc8a1a* (*ncx1*), *ryr2b, atp2a2a (serca2)*] and other factors expressed preferentially in cardiomyocytes (*cx36.7*, *nppa/al/b*; Fig. S1D; Table S1). Not surprisingly, gene ontology (GO) term enrichment analysis uncovered overrepresentation of biological process terms “heart development”, “angiogenesis”, “heart contraction”, “actin cytoskeletal organization”, and “sarcomere organization” in the ALPM-enriched gene set (Fig. S1E, see Table S1 for complete GO term analysis). The successful identification of ALPM markers with well-documented and indispensable roles in cardiopharyngeal progenitors or cardiomyocytes suggests that novel molecular activities of equal importance are likely to be among the uncharacterized ALPM-restricted genes.

**Figure 1.**
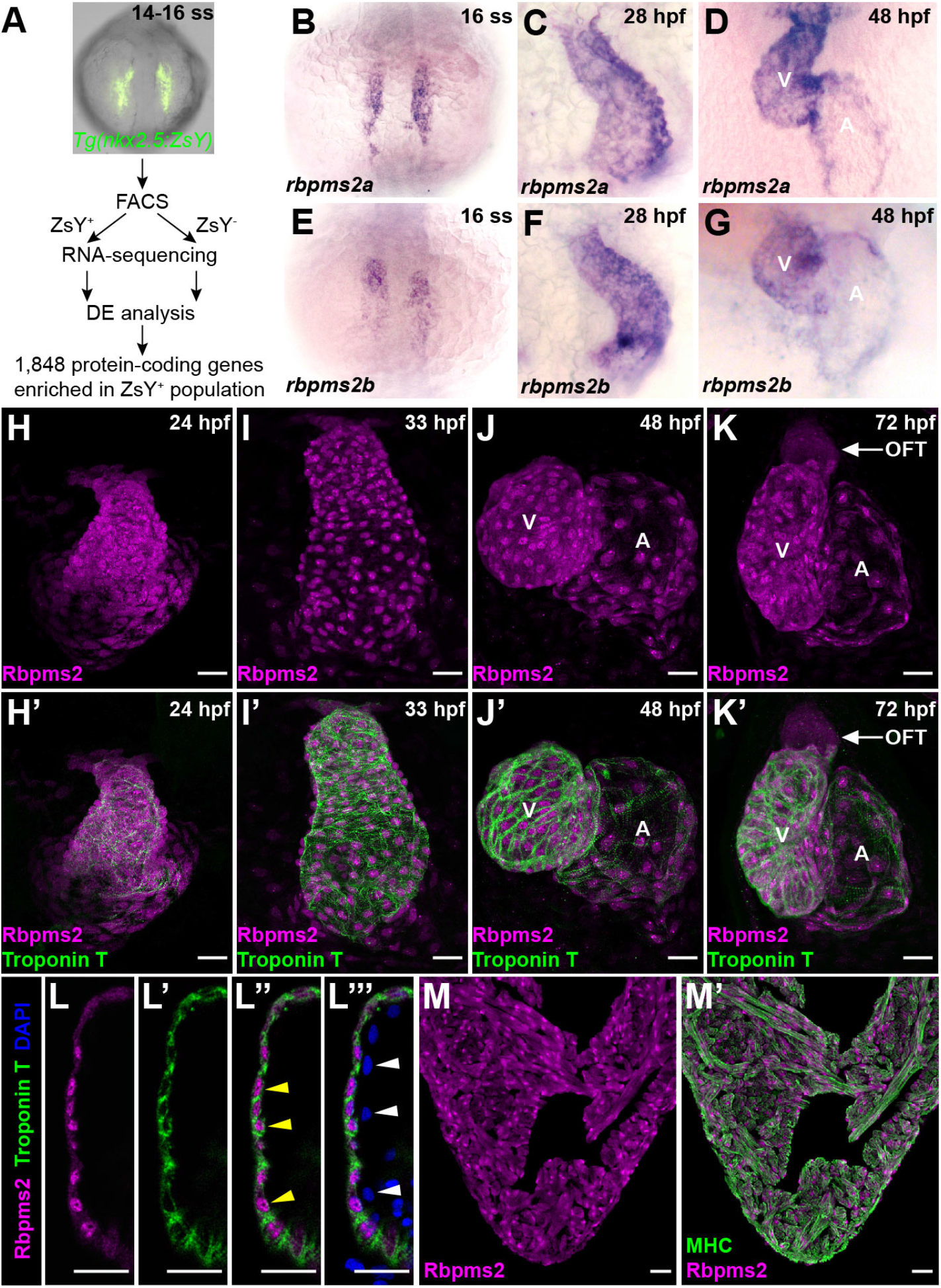
Restricted expression of RNA-binding proteins Rbpms2a and Rbpms2b to the zebrafish myocardium. (A) Workflow used to identify 1848 protein-coding RNAs enriched in ZsYellow-positive (ZsY+) cardiopharyngeal progenitors or cardiomyocytes in the anterior lateral plate mesoderm of 14-16 somite stage (ss) *Tg(nkx2.5:ZsYellow)* embryos. (B-G) Brightfield images of 16 ss (B,E), 28 hours post fertilization (hpf; C,F), and 48 hpf (D,G) embryos processed for in situ hybridization with riboprobes that detect *rbpms2a* (B-D) or *rbpms2b* (E-G). Little to no variation in expression pattern was observed between animals in each group (n>10/group). (H-K’) Confocal projections of hearts in 24 hpf (H,H’), 33 hpf (I,I’), 48 hpf (J,J’), and 72 hpf (K,K’) zebrafish embryos co-immunostained with antibodies that detect Rbpms2 (magenta) or cardiac Troponin T (CT3 antibody; green). Single (H,I,J,K) and merged double channel (H’,I’,J’,K’) images are shown. Little to no variation in expression pattern was observed between animals in each group (n>10/group). (L-L’’’) Single optical section through the ventricular wall of a 48 hpf embryo co-immunostained with antibodies that detect Rbpms2 (magenta) or cardiac Troponin T (CT3 antibody; green) and counterstained with DAPI (blue). Single (L, L’), merged double (L’’), and merged triple (L’’’) channel images are shown. Yellow arrowheads highlight Rbpms2-positive cardiomyocyte nuclei surrounded by Troponin T-positive cytoplasm. White arrowheads highlight Rbpms2-negative nuclei in adjacent endocardial cells lining the ventricular chamber. 4/4 ventricles exhibited this Rbpms2 expression pattern. (M,M’) Confocal projections of a histological section through the ventricle of an adult zebrafish heart co-immunostained with antibodies that detect Rbpms2 (magenta) or Myosin Heavy Chain (MHC; MF20 antibody; green). Single channel (M) and merged double channel (M’) images are shown. 18/18 sections, 6 from each of three hearts, exhibited this Rbpms2 expression pattern. Abbreviations: FACS, fluorescence activated cell sorting, DE, differential expression; V, ventricle; A, atrium; OFT, outflow tract. Scale bars=25μm.

Transcripts encoding <u>R</u>NA <u>b</u>inding <u>p</u>rotein with <u>m</u>ultiple <u>s</u>plicing (variants) 2a (Rbpms2a) and Rbpms2b were enriched 14- and 5-fold, respectively, in the ZsYellow-positive population (Fig. S1D; Table S1), suggesting that these RNA-recognition motif (RRM)-containing proteins localize to the ALPM. Classified as ohnologs, *rbpms2a* and *rbpms2b* encode proteins that are 92% identical and perform redundant functions in oocyte differentiation ^33^. Across the full-length amino acid sequences, Rbpms2a and Rbpms2b share greater than 81% identity with human RBPMS2 (Fig. S2A,B). The shared identity exceeds 92% in the RRMs (Fig. S2A,B). Such a high degree of sequence conservation suggests that the molecular function of RBPMS2 proteins and the properties of their binding site sequences are also likely to be conserved across evolution.

The founding member of the *RBPMS2* gene family, originally termed *hermes*, was discovered in a differential expression screen designed to identify heart-specific transcripts in *Xenopus* ^34,35^. In situ hybridization revealed that *RBPMS2* localizes to the lateral plate mesoderm and embryonic myocardium in both amphibian and avian species ^34,36^. Hearts of embryonic zebrafish were also recently reported to express *rbpms2a* and *rbpms2b* ^33^. Despite the high degree of amino-acid sequence conservation and consistent myocardial-restricted expression pattern across vertebrates, *RBPMS2* has yet to be implicated in any aspect of myocardial biology through genetic loss-of-function studies.

To validate the predicted ALPM localization of *rbpms2a* and *rbpms2b,* we performed in situ hybridization for both genes at the same developmental stage used for profiling *nkx2.5:ZsYellow*-positive ALPM cells (16 ss; Fig. S1). As anticipated, both transcripts localized bilaterally to the ALPM (Fig. 1B,E) in a pattern mirroring early-differentiating cardiomyocytes in at this stage ^37^. Later-stage analysis revealed *rbpms2a* and *rbpms2b* expression in the heart tube at 24-28 hours post fertilization (hpf; Fig. 1C,F; Fig. S3A,C) and in the atrium and ventricle of the two-chambered heart at 48 hpf (Fig. 1D,G; Fig. S3B,D). Consistent with previous reports ^33,38^, extra-cardiac expression was observed in the pronephric duct and retina (Fig. S3A-D)

To investigate the lineage specificity and subcellular localization of Rbpms2a and Rbpms2b in the zebrafish heart, we immunostained wild-type embryos with a monoclonal antibody raised against human RBPMS2, which cross reacts with zebrafish Rbpms2 (see below). Consistent with the distribution of *rbpms2a/b* transcripts, Rbpms2 protein was detected in the heart tube (Fig. 1H,I; Fig. S3E,F), two chambered heart (Fig. 1J,K; Fig. S3G), pronephric duct (Fig. S3E-G) and retina (Fig. S3F,G). Double-immunostaining for Rbpms2 and the myocardial marker cardiac Troponin T (CT3) revealed that Rbpms2 expression localizes to the myocardium at all stages analyzed (Fig. 1H’,I’,J’,K’). Weak expression of Rbpms2 was also evident by 72 hpf in the OFT (i.e. bulbous arteriosus; Fig. 1K,K’). Examination of single optical sections through the ventricular wall, which showed nuclear Rbpms2 signals surrounded by Troponin T-positive cytoplasm, confirmed myocardial localization of Rbpms2, (Fig. 1L-L’’’). Adjacent endocardial cells were devoid of Rbpms2 signals, which highlights the myocardial-specific nature of Rbpms2 expression (Fig. 1L-L’’’). The subcellular distribution of Rbpms2 appeared variable, ranging from a nucleocytoplasmic distribution to exclusively nuclear (Fig. 1H,I,J,K), but the significance of this observation is unknown. Lastly, we documented Rbpms2 expression in the nuclei and cytoplasm of cardiomyocytes in hearts from adult zebrafish (Fig. 1M,M’), suggesting that Rbpms2 functions in the myocardium continuously throughout the lifespan of zebrafish rather than transiently during embryogenesis.

### Generation of Rbpms2-null zebrafish

To discover if *rbpms2a* and *rbpms2b* are required for myocardial development or function, we created mutant alleles of both genes using transcription activator-like effector nucleases (TALENs) ^24^. TALEN pairs were designed to target DNA sequences encoding N-terminal regions of the RRM domains (Fig. S4A,B). We isolated the mutant alleles *rbpms2a^chb3^* and *rbpms2b^chb4^*, which carry frame-shifting deletions and pre-mature stop codons that truncate the proteins by >80% (Fig. 2A,B; Fig. S4A,B). The truncations also remove 82% of the RRMs, suggesting that the RNA-binding activities of the mutant proteins are abolished or significantly compromised. As a result, both alleles are likely to be null. Animals homozygous for either *rbpms2a^chb3^* or *rbpms2b^chb4^* are grossly indistinguishable from siblings during all life stages, which is consistent with a previous study that independently generated mutant alleles of *rbpms2a* and *rbpms2b* ^33^.

**Figure 2.**
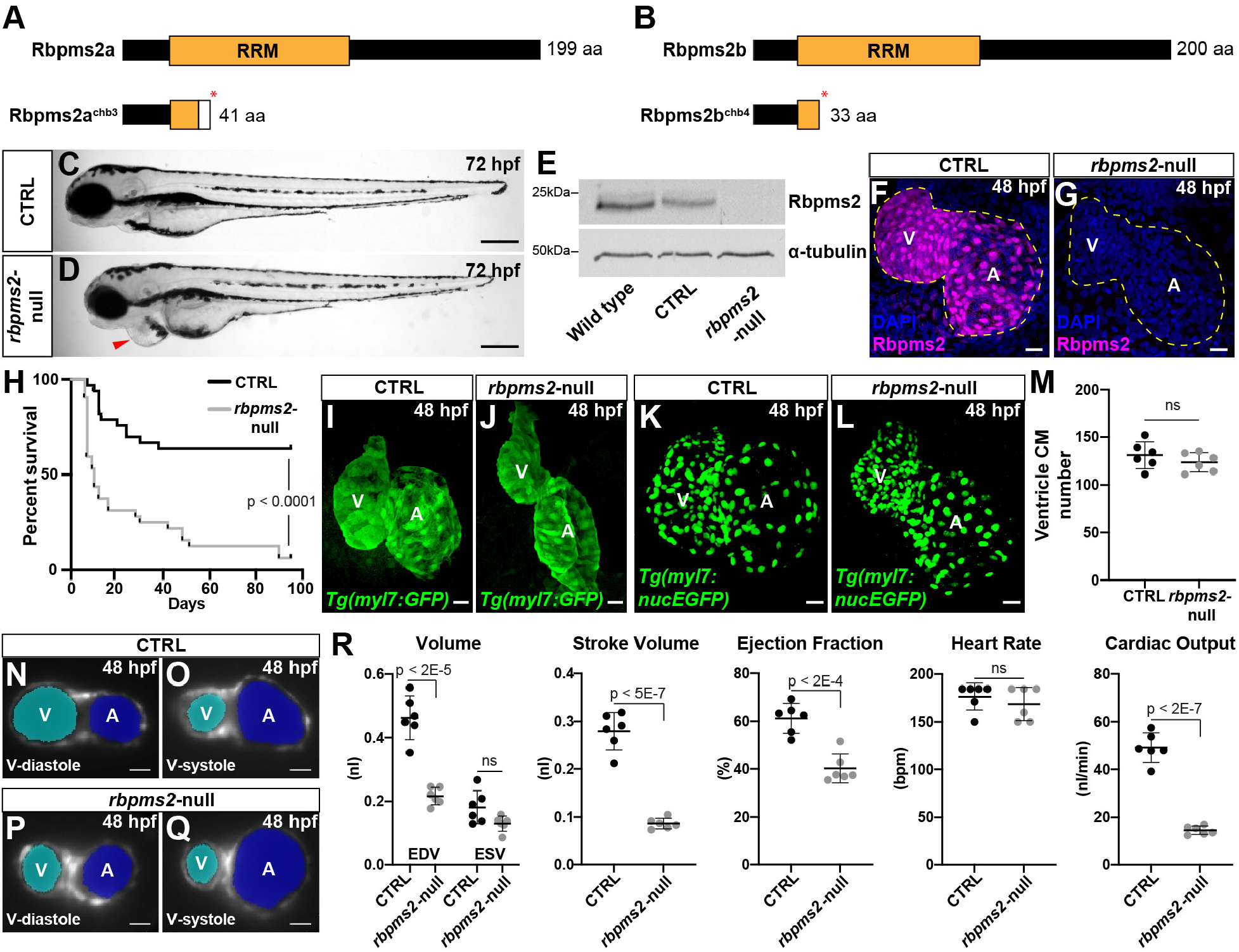
*rbpms2a* and *rbpms2b* are required for ventricular function. (A,B) Schematic diagrams of Rbpms2a and Rbpms2b (top) and the predicted protein products of the *rbpms2a^chb3^* and *rbpms2b^chb4^* alleles (bottom). The asterisks mark the locations of premature stop codons (*) within the RNA-recognition motifs (RRMs) caused by frame-shifting deletions. The white box in (A) shows the location of divergent amino acids prior to the stop codon. (C-D) Brightfield images of control-sibling (C; CTRL) and *rbpms2*-null (*rbpms2a^chb3/chb3^*; *rbpms2b^chb4/chb4^*; D) embryos at 72 hours post fertilization (hpf). Arrowhead highlights pericardial edema in the mutant. Little to no variation in phenotype was observed between animals in each group (n>10/group). Scale bars=250μm. (E) Western blot showing Rbpms2 levels in 48 hpf wild-type (WT), CTRL, and *rbpms2*-null embryos (top) relative to the loading control alpha-tubulin (bottom). (F-G) Confocal projections of hearts in 48 hpf CTRL (F) and *rbpms2*-null (G) embryos immunostained with an antibody that detects Rbpms2 (magenta) and counterstained with DAPI (blue). The dotted lines delineate the myocardium as ascertained from the Rbpms2 signal (F) or *Tg(myl7:eGFP)* reporter signal present in these animals (F,G; not shown). Little to no variation in expression pattern was observed between animals in each group (n>10/group). (H) Kaplan Meier plot showing the survival curves of CTRL (n=33) and *rbpms2*-null (n=33) animals. Statistical significance was determined using a log-rank test. (I-L) Confocal projections of hearts in 48 hpf CTRL (I,K) or *rbpms2*-null (J,L) embryos carrying the *myl7:GFP* (I,J) or *myl7:nucGFP* (K,L) transgenes immunostained with an anti-GFP antibody. For (I,J), little to no variation in phenotype was observed between animals in each group (n>10/group). (M) Dot plot showing ventricular cardiomyocyte (CM) number in CTRL (n=6) and *rbpms2-*null (n=6) hearts at 48 hpf. Error bars show one standard deviation. Statistical significance was determined by an unpaired, two-tailed Student’s t-test assuming equal variances. ns, not significant. (N-Q) Representative single-plane light sheet fluorescence microscopy images of hearts in 48 hpf CTRL (N,O) and *rbpms2*-null (P,Q) embryos carrying the *myl7:GFP* transgene. Images of hearts during ventricular diastole (N,P) or systole (O,Q) are shown with Cardiac Functional Imaging Network (CFIN) overlays of the atrial (blue) and ventricular (cyan) lumens. (R) Dot plots showing end-diastolic volume (EDVs) and end-systolic volume (ESVs), stroke volume, ejection fraction, heart rate, and cardiac output of ventricles in 48 hpf CTRL (n=6) and *rbpms2*-null (n=6) animals measured with CFIN. Error bars show one standard deviation. Statistical significance was determined by an unpaired, two-tailed Student’s t-test assuming equal variances. ns, not significant. Abbreviations: aa, amino acid; V, ventricle; A, atrium. Scale bars=25μm.

To determine if *rbpms2a* and *rbpms2b* perform redundant functions in the heart, we generated and monitored double homozygous embryos [i.e. *rbpms2a^chb3/chb3^; rbpms2b^chb4/chb4^* animals, referred to hereafter as *rbpms2*-null, double-mutant, or double-knockout (DKO) embryos or animals] for cardiac defects. DKO animals are grossly indistinguishable from control siblings for the first ~30 hours of life. By ~33 hpf, they develop a characteristic indentation of the yolk, which is indicative of mild pericardial edema (Fig. S4C,D). At this stage, double-mutants are delayed in initiating blood flow even though peristalsis of the heart tube is readily evident. By 72 hpf, pericardial edema becomes progressively more severe (Fig. 2C,D), suggesting that *rbpms2*-null animals suffer from a cardiac malformation or ongoing functional deficit.

Western blotting and whole mount immunostaining revealed that the zebrafish epitopes recognized by the anti-human RBPMS2 antibody are completely absent in double mutants (Fig. 2E-G), demonstrating that the antibody cross reacts with zebrafish Rbpms2. It also suggests that the mutant mRNAs or proteins are unstable, further suggesting that *rbpms2a^chb3^* and *rbpms2b^chb4^* are bona fide null alleles.

To assess double-mutant viability, we monitored the survival of control-sibling and DKO animals from embryonic stages to adulthood. Over 50% of *rbpms2*-null animals died by two weeks of age (Fig. 2H). Eighty five percent of animals perished by 60 days post fertilization (dpf), and the remaining small percentage of animals did not survive beyond 6 months post fertilization. At all ages, death was generally accompanied by progressively severe pericardial edema, suggesting a cardiac origin. Together, these data reveal that *rbpms2a* and *rbpms2b* ohnologues perform redundant functions that are essential to zebrafish survival.

### rbpms2a and rbpms2b are required for chamber expansion during ventricular filling

To evaluate DKO embryos for structural heart disease, we examined hearts in 48 hpf control-sibling and *rbpms2*-null animals carrying the *myl7:GFP* transgene, which labels the entire myocardium with GFP. Although the two-chambered heart in double mutants displayed normal left-right patterning, the ventricle was displaced anteriorly relative to the atrium (Fig. 2I,J), likely reflecting stretching at the cardiac poles caused by pericardial edema. Atrial size was not obviously altered in *rbpms2*-null animals, but mutant ventricles appeared smaller (Fig. 1I,J). To determine if this might reflect a diminished ventricular cardiomyocyte number, we quantified cardiomyocytes in control-sibling and *rbpms2*-null animals carrying the *myl7:nucGFP* transgene, which labels cardiomyocyte nuclei with GFP (Fig. 2K,L). From this, we learned that ventricular, as well as atrial, cardiomyocyte numbers were unaltered in double mutants (Fig. 2K-M; Fig. S5A), demonstrating that any deviations in chamber size are not attributable to altered myocardial cell content. It also suggests that *rbpms2a* and *rbpms2b* are dispensable for myocardial differentiation from the first and second heart fields, which is the principal method by which cardiomyocytes are produced before 48 hpf ^20,39–41^. These data demonstrate that early cardiac morphogenesis is largely unperturbed in double mutant animals, with the only apparent anatomical abnormality being a reduction in ventricular size.

To characterize potential functional defects in *rbpms2*-null hearts, we measured ejection fraction and cardiac output in 48 hpf animals using our previously described Cardiac Functional Imaging Network [CFIN; ^42^]. To that end, we captured Z-stack dynamic images of GFP+ hearts over several cardiac cycles in control-sibling and DKO *myl7:GFP* transgenic embryos using light sheet fluorescence microscopy. CFIN extracted chamber volumes and identified the maximum and minimum ventricular volumes throughout the cardiac cycle, referred to as end-diastolic volume (EDV; maximum volume during cardiac relaxation) and end-systolic volume (ESV; minimum volume during cardiac contraction), respectively. Whereas the ESV was unchanged in DKO animals, the EDV was significantly reduced (Fig. 2N-R), demonstrating that the ventricle fails to expand during the relaxation phase of the cardiac cycle. As a result, three indices of ventricular function, all of which are calculated from EDV and ESV, were significantly reduced. These include stroke volume (SV), the volume of blood expelled by the ventricle per beat (SV=EDV-ESV), ejection fraction, the percentage of blood expelled by the ventricle per beat (EF=SV/EDV), and cardiac output, the total volume of blood expelled by the ventricle per minute (SV*heart rate) (Fig. 2R). Heart rate was unchanged in the mutant (Fig. 2R). CFIN analysis also revealed an increase in atrial volume (Fig. S5B), which was not grossly obvious after fixation (Fig. 2I,J) and is a common secondary consequence of impaired ventricular function ^43–45^. These data demonstrate that *rbpms2a* and *rbpms2b* are required for ventricular function in the zebrafish embryo. Further, this requirement likely extends into adulthood because the longest-living double-mutant animals exhibited several signs of heart failure such as stunted growth, pericardial edema, and cardiac pathologies including compact, fibrotic ventricles and massively enlarged atria (Fig. S6A-E).

### Identification of differential alternative splicing events in rbpms2-null zebrafish

To identify transcriptomic abnormalities in the hearts of *rbpms2*-null embryos, we performed bulk RNA-sequencing and differential expression analysis on control-sibling and *rbpms2*-null animals at 33 hpf, the earliest developmental stage when DKO animals are distinguishable by morphology (Fig. S4C,D). We reasoned that early-stage analysis would maximize the chances of highlighting primary molecular defects without interference by secondary responses to cardiac dysfunction. Importantly, we confirmed that the primitive ventricle in 33 hpf *rbpms2*-null animals exhibits a functional deficit by documenting reduced myocardial wall movement at the arterial pole of the heart tube (Fig. S7A). We also confirmed that total cardiomyocyte numbers are unaltered at 33 hpf in double mutants (Fig. S7B), reducing the chances that any transcriptomic abnormalities would be attributable to altered cellular composition. Lastly, we profiled whole embryos, instead of purified hearts, because effective protocols for isolating hearts or cardiomyocytes at this early stage do not currently exist. Nonetheless, because *rbpms2a* and *rbpms2b* are expressed with exquisite specificity in the heart, retina and pronephric duct (Fig. S3F), we anticipated that the transcriptional changes occurring in the myocardium would be a discernable subset of those occurring in the whole animal.

While RNA-sequencing and gene-level differential expression analysis highlighted only 2 downregulated transcripts in *rbpms2*-null animals (|FC|>1.5; p-adj<0.05; Table S2; Figure S8A), isoform-level differential expression analysis yielded 78 downregulated and 74 upregulated transcripts (|FC|>1.5; p-adj<0.05; Table S3; Fig. S8B). GO term analysis did not identify overrepresentation of any terms in either gene set [False Discovery Fate (FDR)<0.05]. However, among the downregulated transcripts, there were three well-characterized myocardial genes including *phospholamban* (*pln*; FC=−43.00; p-adj=1.53E-09), *atrial myosin heavy chain 6* (*myh6*; FC=−2.17; p-adj=1.52E-11), and *natriuretic peptide precursor a* (*nppa*; FC=-1.57; p-adj=0.019; Fig. S8B; Table S2). We confirmed downregulation of *nppa* by reverse-transcription quantitative PCR (RT-qPCR) at 33hpf (Fig. S7C). However, this reduction was short lived because two subsequent developmental stages showed increased *nppa* expression (Fig. S7C), which is characteristic behavior for *nppa* in response to cardiac stress ^46^. RT-qPCR analysis of *myh6* revealed a downward trending but statistically insignificant change in double mutants (Fig. S7D). Ultimately, downregulation of either gene is unlikely to explain the ventricular phenotype in *rbpms2*-null mutants because *nppa-*null zebrafish do not exhibit any gross cardiac defects ^47^ and the primary defect in *myh6*-deficient animals is a silent atrium ^48^.

Of the two *pln* genes in the zebrafish genome (GRCZ11), *si:ch211-270g19.5* (ENSDARG00000097256) and *pln* (ENSDARG00000069404), the latter is alternatively spliced according to publicly available cDNA sequencing data. Interestingly, only one of the alternatively-spliced *pln* isoforms (ENSDART00000141159; *pln-203*) was found to be downregulated in *rbpms2*-null animals (FC=-43.00; p-adj1.53E-09; Fig. S8B; Table S2). The other *pln* transcript (ENSDART00000133911;*pln-202*) was unchanged (FC=1.37; p-adj>0.99; Table S2), which highlighted the possibility that Rbpms2 functions in the zebrafish myocardium to regulate alternative splicing, a molecular function previously ascribed to the closely related RBPMS protein ^49^, but not to RBPMS2 itself ^35^.

To investigate this possibility directly, we performed replicate multivariate analysis of transcript splicing (rMATS) ^50^ on the same RNA-sequencing dataset and identified 150 differential alternative splicing events (ASEs) in *rbpms2*-null animals (0 uncalled replicates; FDR<0.01; |Incleveldifference|>0.1; Fig. 3A,B; Table S3). The most common ASE was skipped exon (SE), representing over 40% of the total ASEs, followed by alternative 3’ splice site (A3SS, 23%), alternative 5’ splice site (A5SS, 21%), mutually exclusive exons (MXE, 10%), and retained intron (RI, 6%) (Fig 3A,C). Gene ontology (GO) term enrichment analysis did not identify overrepresentation of any terms among the differentially spliced genes. However, a limited number of myocardial genes were found to be differentially spliced in *rbpms2-*null animals including *myosin binding protein C3* (*mybpc3*) and *myomesin 2a* (*myom2a*), both of which encode sarcomere components ^51–54^, and *solute carrier family 8 member 1a* [*slc8a1a*; also termed the *sodium calcium exchanger 1* (*ncx1*)] which encodes a plasma membrane transporter responsible for extruding calcium from cardiomyocytes during cardiac relaxation ^55,56^.

**Figure 3.**
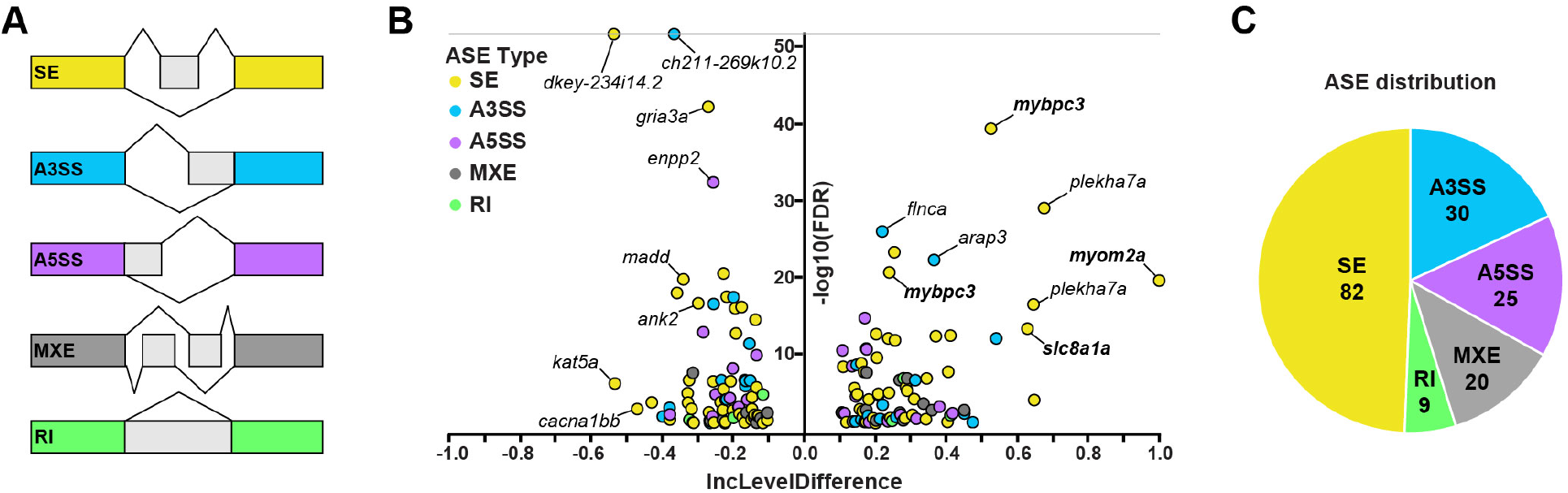
Identification of differential alternative splicing events in *rbpms2*-null embryos. (A) Schematic diagrams of the categories of alternative splicing events (ASEs) investigated by the RNA-seq Multivariate Analysis of Transcript Splicing (rMATS) algorithm including skipped exon (SE), alternative 3’ splice site (A3SS), alternative 5’ splice site (A5SS), mutually exclusive exon (MXE), and retained intron (RI). (B) Volcano plot showing the inclusion (Inc) level differences (control-sibling Inclevel - *rbpms2*-null Inclevel) and False Discovery Rates (FDR) for differential ASEs between 33 hours post fertilization (hpf) control-sibling and *rbpms2*-null animals (0 uncalled replicates; FDR<0.01; |Incleveldifference|>0.1;). The ASEs shown in bold were validated with splicing-sensitive qPCR and in situ hybridization. (C) Pie chart showing the proportions of differential ASEs in each category in *rbpms2*-null animals.

In the case of *mybpc3*, exons 2 and exons 10 were identified as cassette exons whose inclusion levels (Inclevels) in mature *mybpc3* transcripts were found to be significantly lower in *rbpms2*-null animals (exon2 lnclevel=0.031; exon 10 Inclevel=0.057) when compared to those in control siblings (exon2 lnclevel=0.557; exon10 lnclevel=0.296; exon 2 lncleveldifference=0.526, FDR=4.77E-40; exon 10 lncleveldifference=0.24, FDR=2.38E-21; Fig. 3B; Fig.4A; Table S4). The same was true for exon 16 of *myom2a* (control lnclevel=1, DKO lnclevel=0, lncleveldifference=1, FDR 2.74E-20; Fig. 3B; Fig.4A; Table S4) and exon 5 of *slc8a1a* (control lnclevel=1, DKO lnclevel=0.37, lncleveldifference=0.629, FDR=4.92E-14; Fig. 3B; Fig.4A; Table S4). In all cases, exon inclusion was higher in in control siblings, which means that these exon sequences were significantly reduced or absent in *rbpms2*-null animals.

**Figure 4.**
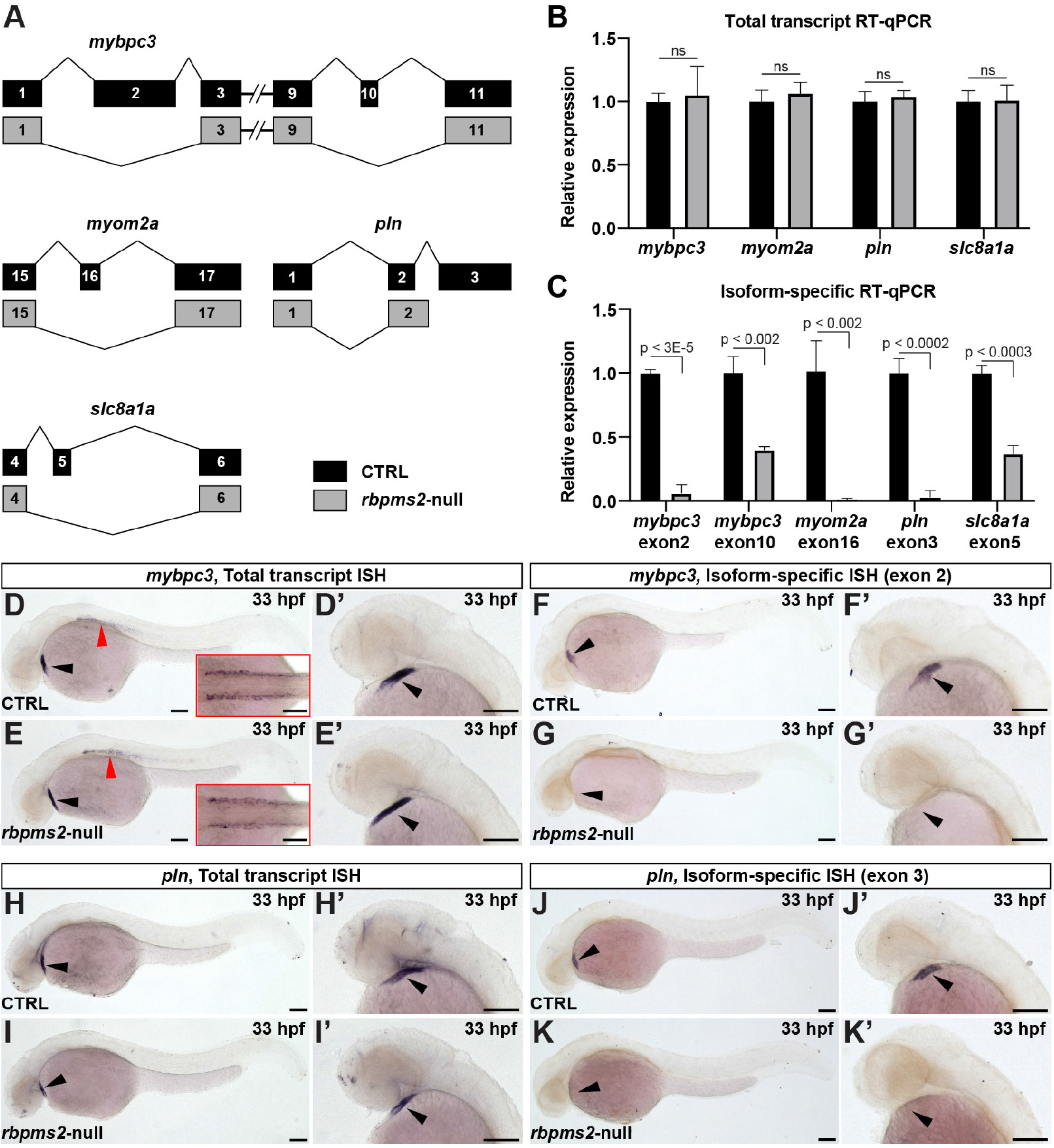
Validation of differential alternative splicing events in *rbpms2*-null animals. (A) Schematic diagrams of alternative splicing events (ASEs) identified in control-sibling (CTRL) or *rbpms2*-null animals at 33 hours post fertilization (hpf) by the RNA-seq Multivariate Analysis of Transcript Splicing (rMATS) or mixture of isoforms (MISO) algorithms. The legend indicates that the isoforms shown on top (black) are significantly reduced or absent in *rbpms2*-null animals and replaced by the isoforms shown on the bottom (gray). (B) Bar graph showing the relative expression levels of the four genes shown in (A), independent of the alternatively spliced exons, between 48 hours post fertilization control-sibling and DKO animals. (C) Bar graph showing the relative expression levels of the indicated, exon-containing isoforms [(shown in black in (A)] of the same four genes between control-sibling and *rbpms2*-null animals at 48 hpf. For (B,C), error bars show one standard deviation. Statistical significance was determined by an unpaired, two-tailed Student’s t-test assuming equal variances. ns, not significant. (D-K’) Brightfield images of 33 hpf CTRL (D,D’,F,F’,H,H’J,J’) and *rbpms2*-null (E,E’,G,G’,I,I’,K,K’) embryos processed for in situ hybridization with riboprobes that detect total pools (D-E’; H-I’) or specific exon-containing mRNA isoforms (F-G’; J-K’) of *mybpc3* (D-G’) or *pln* (H-K’). The cardiac regions in (D,E,F,G,H,I,J,K) are shown at higher zoom in (D’,E’,F’,G’,H’,I’,J’,K’). Red arrowheads in (D,E) highlight expression of *mybpc3* in skeletal muscle, which is shown from a dorsal view in the insets. Black arrowheads in (D-K) indicate the linear heart tube. Little to no variation in expression pattern was observed between animals in each group (n>10/group). Scale bars=100μm.

Despite supporting evidence from differential isoform expression analysis (Fig. S8B; Table S2), rMATS did not identify alternative splicing of *pln*, likely because it involves a terminal exon (Fig. 4A), which rMATS is not designed to evaluate (Fig. 3A). Therefore, we employed the mixture of isoforms (MISO) algorithm ^57^ to evaluate alternative splicing of *pln* in *rbpms2*-null animals and found strong statistical evidence that the exon 3-containing *pln*-203 isoform (ENSDART00000141159) is uniquely reduced in mutant animals. Specifically, the Bayes factor was largely above the accepted cutoff of 5 for differences in ENSDART00000141159 versus ENSDART00000133911 usage in WT and double-mutant animals across all five replicate pairs (Table S5), which is strongly suggestive of differential alternative splicing. This conclusion was also supported by genotype-specific differences in read densities across the locus (Fig. S9). Collectively, these data demonstrate that *rbpms2*-null embryos display abnormal splicing of genes that are essential to cardiac function.

### Validation of differential alternative splicing events in rbpms2-null zebrafish

Next, we used splicing-sensitive RT-qPCR to validate differential alternative splicing of *mybpc3*, *myom2a*, *slc8a1*, and *pln* in *rbpms2*-null animals at 48 hpf (Fig. 4A). To that end, we designed two primer pairs for each transcript. One was designed to measure relative “total transcript” levels between sibling control and *rbpms2*-null animals, independent of the alternatively spliced exon. The other was designed to quantify relative “isoform-specific” levels by having one of the primer-binding sites contained within the exon of reduced inclusion according to rMATS or MISO (Fig. 4A). For all four genes, total transcript levels were not significantly altered in *rbpms*2-null animals (Fig. 4B). However, the specific exon-containing isoforms were reduced by at least 50% in all cases, and by greater than 95% in three cases, indicative of almost-complete exon exclusion (Fig. 4C). Taken together, these data provide strong evidence that Rbpms2 activity promotes the inclusion of specific exons into mature cardiac transcripts. Importantly, we verified that *mybpc3* and *pln* are also abnormally spliced earlier in development (33 hpf), when signs of cardiac distress first become visible in double mutants, reducing the likelihood that abnormal splicing is a secondary consequence of cardiac dysfunction (Fig. S7E,F).

To confirm that the DKO splicing abnormalities localize to the myocardium, we performed splicing-sensitive in situ hybridization analysis for two genes, *mybpc3* and *pln*, with *rbpms2*-dependent exons large enough to generate effective exon-specific anti-sense riboprobes (i.e. exon 2 of *mybpc3* and exon 3 of *pln*; Fig. 4A). In a strategy analogous to the splicing-sensitive qPCR, we designed a pair of riboprobes for each gene. One probe detected the total transcript pools of *mybpc3* or *pln*, independent of the alternatively spliced exons (“Total-transcript” in situ hybridization). The other probe detected the alternatively spliced exon sequences exclusively. In control-sibling animals at 33 hpf, total *mybpc3* transcripts were localized to both cardiac and skeletal muscle (Fig. 4D,D’), and total *pln* transcripts were detected exclusively in the heart (Fig. 4H,H’). These expression patterns were unaffected by the loss of *rbpms2* (Fig. 4E,E’,I,I’), which is consistent with the total transcript qPCR analysis (Fig. 4B). Isoform-specific in situ hybridization analysis revealed expression of the *rbpms2*-dependent *mybpc3* and *pln* exons exclusively in the hearts of control-sibling animals (Fig. 4F,F’,J,J’). Consistent with the isoform-specific qPCR analysis (Fig. 4C), double-mutant animals were completely devoid of these exon sequences (Fig. 4G,G’,K,K’). Taken together, these data implicate *rbpms2* as a positive regulator of exon inclusion in the zebrafish embryonic myocardium.

For those alternative splicing events validated by qPCR or in situ hybridization (*mybpc3*, *myom2a*, *pln*, and *slc8a1a*), exon exclusion in mutant hearts maintains the open reading frame, indicating that the hearts of mutant animals express proteins lacking amino acid sequences encoded by the alternatively spliced exons (Fig. 4E,E’,G,G’I,I’,K,K’). In the case of *mybpc3,* exon 2 encodes a domain termed C0, which is contained exclusively within the cardiac isoform of Myosin Binding Protein C in higher vertebrates (MYBPC3) but is absent from the skeletal muscle isoform [MYBPC2; ^58^]. Similarly, *mybpc3*’s exon 10 encodes cardiac-specific amino acids that become phosphorylated by protein kinase A (PKA) in response to *β*-adrenergic signaling ^59,60^. For Myom2a, the short stretch of amino acids that disappears in DKO animals is of unknown function. For *pln*, excluding exon 3 shortens the protein by 5 amino acids on the C-terminus, which has been shown to increase SERCA inhibitory activity *in vitro* ^6^. Lastly, the *slc8a1a* exon with reduced inclusion in DKO animals is one of six conserved exons that are alternatively spliced in higher vertebrates to control the Ca2+ sensitivity of the transporter ^61,62^.

### Zebrafish rbpms2-null ventricular cardiomyocytes exhibit myofibrillar disarray and calcium handling defects

Because Mybpc3 and Myom2a are sarcomere components, we evaluated 48 hpf control-sibling and double-mutant hearts for myofibrillar defects after immunostaining with antibodies that recognize thin filaments, thick filaments, or Z-disks. For all epitopes, prominent striations were apparent in control-sibling animals reflecting the tandem arrays of sarcomeres composing myofibrils, which themselves are transversely aligned and bundled near the cell cortices (Fig. 5A,A’,C-C”,E). By contrast, despite the readily-evident detection of epitopes in DKO hearts, striations were significantly reduced or absent because the myofibrils were in a state of disarray (Fig. 5B,B’,D-D”,F). We confirmed that myofibril disarray was also present in ventricular cardiomyocytes of double mutant animals at an earlier developmental stage of 33 hpf (Fig. S7G-H’). Lastly, this defect is specific to ventricular cardiomyocytes because myofibril organization in atrial cardiomyocytes of control and DKO embryos was grossly indistinguishable (Fig. S7I-J’). Taken together, these data demonstrate that Rbpms2 activity is required for establishing or maintaining the three-dimensional architecture of myofibrils in ventricular cardiomyocytes of the zebrafish embryo.

**Figure 5.**
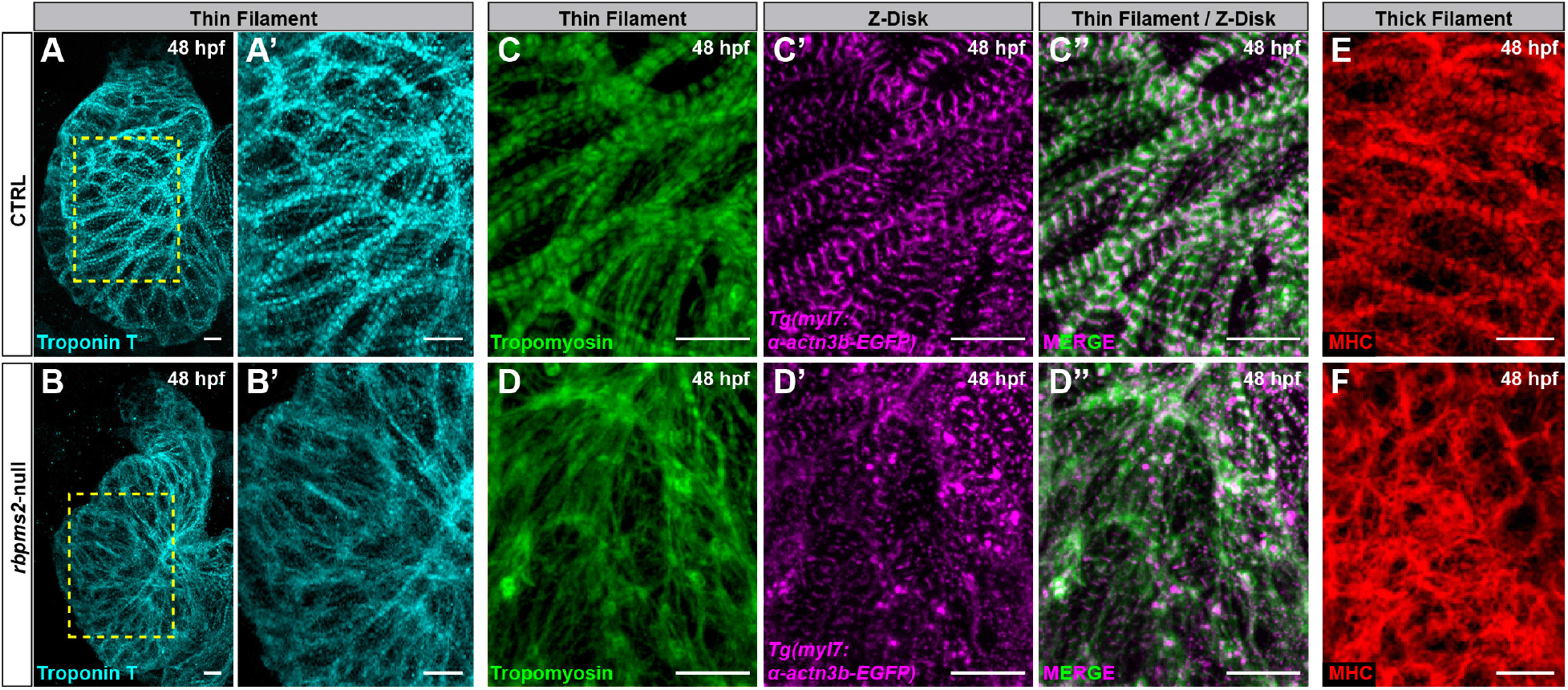
Myofibrillar disarray in *rbpms2*-null ventricular cardiomyocytes. (A-B’) Confocal projections of ventricles in 48 hours post fertilization (hpf) control-sibling (CTRL) or *rbpms2*-null embryos immunostained with an anti-TroponinT antibody (CT3) to visualize thin filaments. Regions demarcated in (A) and (B) are enlarged in (A’) and (B’). (C-D’’) Confocal projections of ventricles in 48 hpf CTRL and *rbpms2*-null *Tg(α-actn3b-EGFP)* animals co-immunostained with antibodies that detect Tropomyosin (CH1; green) to visualize thin filaments or GFP (magenta) to visualize Z-disks. (E-F) Confocal projections of ventricles in 48 hpf CTRL and *rbpms2*-null embryos immunostained with an antibody that detects myosin heavy chain (MF20) to visualize thick filaments. Little to no variation in protein distributions were observed between animals in each group (n>10 animals/group). Scale bars=10μm.

Lastly, because *pln* and *slc8a1a* regulate calcium handing in cardiomyocytes ^56,63–65^, we evaluated *rbpms2*-null hearts for abnormalities in the amplitude or duration of cytosolic calcium transients, which stimulate cardiomyocyte contraction. To that end, control-sibling and *rbpms2*-null hearts at 48 hpf were explanted and loaded with the dual-wavelength calcium-sensitive dye Fura-2. High-speed imaging was performed to obtain a ratiometric indicator of fluctuating calcium in cardiomyocytes over several cardiac cycles ^66^. From this, we measured the duration and amplitude of the calcium transients in the atrium and ventricle of control-sibling and *rbpms2*-null hearts. Whereas these values were not significantly different in the atrium of double mutant hearts (Fig. 6A,B,D-F,H), both the duration and amplitude of the calcium transients in double mutant ventricles were reduced in the mutant ventricle (Fig. 6A-C; E-G), demonstrating that Rbpms2 is indispensable for the regulation of calcium handling in ventricular cardiomyocytes of the zebrafish embryo.

**Figure 6.**
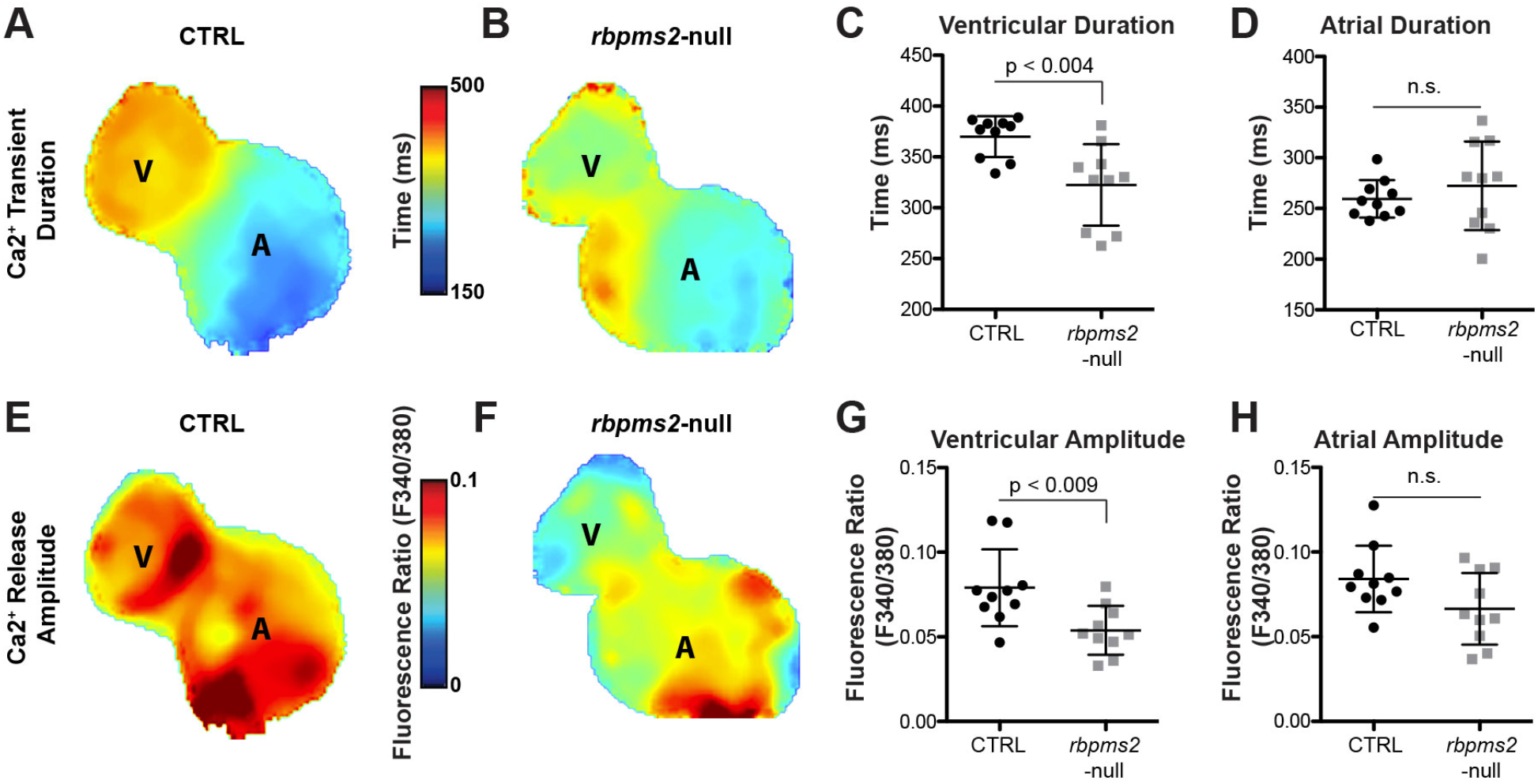
*rbpms2*-null ventricular cardiomyocytes exhibit reductions in Ca2+-transient duration and amplitude. Color maps (A,B,E,F) and dot plots (C,D,G,H; n=10/group) of Ca2+-transient duration (A-D) and Ca2+ release amplitude (E-F) in the ventricle (A-C; E-G) and atrium (A,B,D,E,F,H) of 48 hours post fertilization control-sibling (CTRL; A,C,D,E,G,H) and *rbpms2*-null (B-D; F-H) hearts. Color code depicts localized Ca2+-transient duration in milliseconds (ms; A,B) or the ratiometric fluorescence indicator of Ca2+-release amplitude. Each dot represents center-chamber measurements from one heart. Error bars show one standard deviation. Statistical significance was determined by an unpaired, two-tailed Student’s t-test assuming equal variances. ns, not significant.

### Human RBPMS2-null cardiomyocytes exhibit myofibrillar disarray and calcium handling defects

To determine if the cellular function of *RBPMS2* is conserved in human cardiomyocytes, we targeted *RBPMS2* in human induced pluripotent stem cells (hiPSCs) using CRISPR/Cas9-mediated genome editing and analyzed phenotypes that arose following differentiation into hiPSC-derived cardiomyocytes (hiPSC-CMs). We targeted the DNA sequence encoding RBPMS2’s RNA recognition motif and isolated a clone with a biallelic 2 base-pair deletion (Fig. S10A; *ΔRBPMS*), which truncates both the protein and the RRM by ~70% due to a premature stop codon (Fig. 7A; Fig. S10B). Quantitative RT-PCR analysis revealed a significant 82% reduction in *RBPMS2* transcripts in *ΔRBPMS* hiPSCs (Fig. 7B) and hiPSC-CMs (Fig. 7C), suggesting that the mutant transcripts are susceptible to non-sense mediated decay. This observation, in combination with the large size of the C-terminal truncation (Fig. 7A) strongly suggest that *ΔRBPMS2* cells are null for *RBPMS2*.

**Figure 7.**
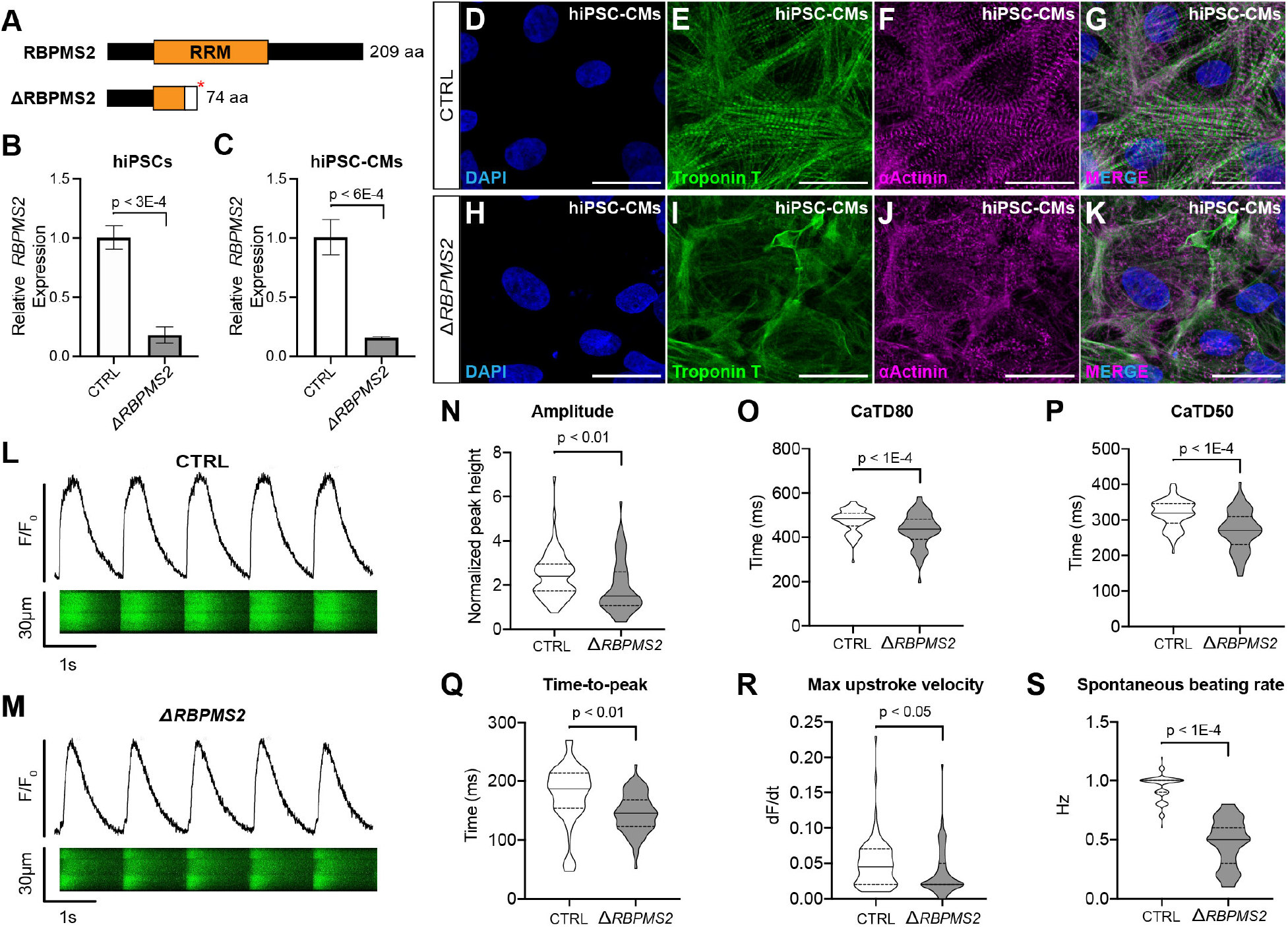
Human *RBPMS2*-null cardiomyocytes exhibit myofibrillar disarray and Ca2+ handling defects similar to those in zebrafish *rbpms2*-null cardiomyocytes. (A,B) Schematic diagrams of human RBPMS2 (top) and the predicted protein product of the *ΔRBPMS2* null allele created with CRISPR/Cas9-mediated genome editing (bottom). The asterisk shows the location of a premature stop codon within the RNA-recognition motif (RRM) caused by a frame-shifting 2 base pair deletion. The white box shows the location of divergent amino acids prior to the stop codon. (B,C) Bar graphs showing the relative expression levels of *RBPMS2* in parental control (CTRL) and *ΔRBPMS2* human induced pluripotent stem cells (hiPSCs; B) and cardiomyocytes (hiPSC-CMs; C) after 15 days of directed differentiation. Error bars show one standard deviation. Statistical significance was determined by an unpaired, two-tailed Student’s t-test assuming equal variances. (D-K) Representative confocal projections of CTRL (D-G) and *ΔRBPMS2* (H-K) hiPSC-CMs immunostained with antibodies that detect TroponinT (CT3; green) or Alpha Actinin (anti-Sarcomeric Alpha Actinin; magenta) to visualize thin filaments or Z-disks, respectively, and counterstained with DAPI (blue). Single (D-F; H-J) and merged triple (G,K) channel images are shown. Little to no variation in staining pattern was observed between cells in each experimental group (n>50 cells/group from at least three wells from two separate differentiations were examined). (L,M) Representative traces of fluorescence intensity over baseline (F/F_o_; top) derived from line scans (bottom) of CTRL (L) and *ΔRBPMS2* (M) hiPSC-CMs loaded with fluo-4 and paced at 1Hz. (N-S) Violin plots showing Ca2+-transient amplitude (N), duration at 80% (O) and 50% (P) repolarization, time to peak amplitude (Q), upstroke velocity (R), and un-paced spontaneous beating rate (S). Statistical significance was determined by an unpaired, two-tailed Student’s t-test assuming equal variances. n=80 cells/group, 40 each from two separate differentiations. Scale bars=25μm.

To determine if *RBPMS2*-null human cardiomyocytes exhibit cellular phenotypes similar to those observed in ventricular cardiomyocytes from *rbpms2*-null zebrafish (Fig. 5A-F), we analyzed myofibril structure in control-parental and *ΔRBPMS2* hiPSC-CMs by double immunostaining with antibodies that detect thin filaments or Z-disks. The hallmark striations created by thin filaments, which were readily apparent in control cells (Fig. 7D,E,G), were almost completely absent in *ΔRBPMS2* hiPSC-CMs (Fig. 7H,I,K). Moreover, whereas the Z-disks in control cardiomyocytes were prominent, well defined and arrayed in parallel (Fig. 7D,F,G), those in *ΔRBPMS2* hiPSC-CMs were smaller, punctate, and disorganized (Fig. 7H,J,K). Taken together, these data uncover a conserved role for RBPMS2 in establishing or maintaining myofibril organization in human cardiomyocytes.

Lastly, we evaluated control-parental and *ΔRBPMS2* hiPSC-CMs for defects in calcium handling. hiPSC-CMs cells were loaded with the calcium-sensitive dye Fluo-4, paced at 1Hz and imaged by line scanning confocal microscopy to capture intracellular Ca2+ dynamics (Fig. 7L,M). Similar to ventricular cardiomyocytes from *rbpms2*-null zebrafish, *ΔRBPMS2* hiPSC-CMs exhibited significant decreases in Ca2+-transient amplitude (Fig. 7N) and transient duration (Fig. 7O,P). These phenotypes were accompanied by decreases in time-to-peak (Fig. 7Q) and maximum upstroke velocity (Fig. 7R). Lastly, without pacing, *ΔRBPMS2* hiPSC-CMs contracted at a lower spontaneous beat frequency (Fig. 7S). These data demonstrate that human cardiomyocytes rely on RBPMS2 function for optimal calcium handling. Taken together, the overlap in sarcomere and calcium handling phenotypes observed between *Rbpms2*-null zebrafish and *ΔRBPMS2* hiPSC-CMs suggest that the cellular function(s) of this myocardial RNA-binding protein is conserved in human cardiomyocytes.

## DISCUSSION

Our study identifies Rbpms2 proteins as critical RNA splicing factors that regulate tissue-specific isoform expression within the ventricular myocardium. Specifically, we show that zebrafish Rbpms2a and Rbpms2b are redundantly required for ventricular form and function in embryonic zebrafish. We then show that Rbpms2 proteins function to promote the generation of distinct isoforms within the heart, including a previously undocumented cardiac-specific isoform of *mybpc3*. Further, we found that *rbpms2*-null hearts have disorganized sarcomeres and disrupted calcium handling, which is phenotypically consistent with the broad classes of alternatively spliced transcripts identified by our RNA-seq. Lastly, we discovered that RBPMS2-deficient human iPSC-CMs exhibit defects in sarcomere structure and calcium handling that are strikingly similar to what we observed in zebrafish ventricles.

Our characterization of Rbpms2 provides further evidence that RNA splicing within the heart may be a much more pervasive and physiologically important mode of gene regulation than previously appreciated. While our results reveal a clear connection between Rbpms2 and RNA splicing, these proteins may function in additional capacities within tissues outside the heart. Previous reports in zebrafish have shown that Rbpms2 proteins contribute to the establishment of oocyte polarity and development by influencing the localization of RNAs such as *bucky ball* (*buc*) to the Balbiani body^33,67,68^. Further research in xenopus and chicken have described similar roles for Rbpms2 in RNA stability and localization within intestinal smooth muscle, kidney cells, and retinal ganglion cells^38,69,70^. Interestingly, the localization of Rbpms2 to stress granules has been observed in several of these models^69,71^, indicating that this function may be conserved in multiple tissues. It will be important to determine if RBPMS2 functions in this capacity within the heart, or whether this is a tissue-specific function that is restricted to other RBPMS2-positive organs such as the eye and nephron.

Even though we found that Rbpms2 regulates the splicing of genes whose orthologs are known to cause cardiomyopathy, it is unclear whether mutations in RBPMS2 segregate with human disease. Our data therefore warrants a careful re-evaluation of existing GWAS datasets and provides new incentives to examine the RBPMS2 locus in CHD patients. However, given the severity of cardiac phenotypes observed in our loss-of-function experiments, it remains possible that embryonic lethality may confound GWAS results. Additionally, the upstream regulators of Rbpms2 remain unknown. Further studies will be necessary to identify these upstream factors in order to determine if they contribute to disease states through a more subtle modulation of Rbpms2 function. Given that Rbpms2 is itself alternatively spliced in vertebrates, it is also possible that its function may be regulated by yet another upstream splicing factor. Collectively these studies will evaluate RBPMS2 function within the broader context of human cardiomyopathy and heart failure.

## ACKNOWLEDGEMENTS

Fluorescence activated cell sorting was performed at the Harvard Stem Cell Institute-Center for Regenerative Medicine Flow Cytometry Core. Customized TALENs were generated by the Broad Institute Genetic Perturbation Platform (Director: John Doench, PhD). Library preparation, RNA-sequencing, and bioinformatics analyses were performed at the MIT BioMicroCenter (Director: Stuart Levine, PhD). Ryan Abo of the MIT BioMicroCenter provided assistance with bioinformatics analysis. Sanger sequencing and DNA fragment analysis were performed at the Massachusetts General Hospital Computational and Integrative Biology DNA Core (Director: Amy Avery).

## FUNDING

AAA was supported by an American Heart Association Postdoctoral Fellowship (20POST35110027). AAA and AS were supported by a National Institutes of Health (NIH) training grant awarded to Massachusetts General Hospital (MGH) (T32HL007208; PI: A. Rosenzweig). The Burns Laboratory was supported by NIH grants 7R01HL139806 (formerly 1R01HL139806) and 7R35HL135831 (formerly 1R35H135831) to CGB and CEB, respectively, by awards from the Executive Committee on Research at MGH to CGB, and by Department of Defense Peer-Reviewed Medical Research Program (PRMRP) grant PR190628 to CEB. CEB was a d’Arbelhoff MGH Research Scholar. CGB was a Hassenfeld Cardiovascular Scholar at MGH. The Burns Laboratory was supported, in part, by funds from the Boston Children’s Hospital Department of Cardiology.

MT was supported by an NIH training grant awarded to Boston Children’s Hospital (T32HL007572; PI: W. Pu). The bioinformatics analysis was supported in part by the Koch Institute Support (core) Grant P30-CA14051 from the National Cancer Institute.

## COMPETING INTERESTS

The authors declare no competing interests.

## Notes

### Competing Interest Statement

The authors have declared no competing interest.

