## Supplemental information for "RBPMS2 is a conserved regulator of alternative splicing that promotes myofibrillar organization and optimal calcium handling in cardiomyocytes"

### EXPANDED MATERIALS & METHODS

#### Sample preparation for RNA-sequencing and differential expression analysis

*Tg(nkx2.5:ZsYellow)* embryos at the 14-16 somites stage were dechorionated with pronase and washed several times with excess E3 medium. Approximately 1,200 embryos were transferred to a 1.5ml Eppendorf tube, centrifuged at low speed, and washed three times with 0.9X PBS. Embryos were suspended in 750μl of ice-cold 0.9X PBS plus 5% FBS (PBS/FBS) and homogenized with a disposable plastic pestle before being drawn up-and-down multiple times into a p1000 micropipette. The cells were passed over a 40μm nylon cell strainer and collected in a pre-chilled 50ml Falcon tube on ice. The strainer was subsequently rinsed with 25ml of ice-cold PBS/FBS and the flowthrough was collected in the same Falcon tube. Cells were pelleted by low-speed centrifugation for 5 minutes at 4°C. The majority of supernatant was removed with a Pasteur pipette and the pellet was resuspended in 2ml of ice-cold PBS/FBS. Cells were passed over another 40μm strainer, collected in a pre-chilled 15ml Falcon tube and pelleted by low-speed centrifugation for 5 minutes at 4°C. Cells were resuspended in approximately 1ml ice-cold PBS/FBS and stored on ice. Fluorescence activated cell sorting was used to separate ZsYellow-positive from ZsYellow-negative cells. Both subpopulations were collected independently and frozen. From each sort, averages of 39K ZsYellow-positive cells (range 23-68K) and 50K ZsYellow-negatieve cells were collected. Total RNA was purified using the RNeasy Kit (Qiagen Sciences Inc). Material from either 4 (ZsYellow-positive) or 5 (ZsYellow-negative) sorts was combined to generate single replicates. Two replicates per experimental group, consisting of RNA purified from 147K or 166K total ZsYellow-positive cells and 250K total ZsYellow-negative cells, were included in the RNA-sequencing and differential expression analysis.

Control-sibling and *rbpms2*-null embryos at 33 hpf were separated based on morphology and collected into pools (10 animals per pool). Pools were mechanically homogenized by vigorous pipetting in 500µl TRIzol Reagent (Thermo Fisher Scientific). For each pool, a 200µl sample of the homogenization solution was removed and phase-separated to obtain DNA for confirmatory genotyping. The remaining 300µl was processed via Direct-zol RNA MicroPrep columns (Zymo Research) to obtain total RNA for one biological replicate. Five biological replicates for each experimental group were included in the RNA-sequencing and differential expression analysis.

#### *Tg(nkx2.5:ZsYellow)* FACS RNA-Seq strategy and analysis

RNA sample quality was evaluated using a 2100 Bioanalyzer Instrument (Agilent Technologies). Approximately 50ng of RNA per sample were used to prepare sequencing libraries with the Clontech low-input RNA library preparation kit. Libraries were sequenced as paired-end 2x50nt on an Illumina HiSeq2500 instrument. Quality control was performed by aligning reads against Zv9/danRer7 using bwa mem v. 0.7.12-r1039 with flags –t 16 –f and mapping rates, fraction of multiply-mapping reads, number of unique 20-mers at the 5' end of the reads, insert size distributions and fraction of ribosomal RNAs were calculated using dedicated perl scripts and bedtools v. 2.25.0.64 [72]. In addition, each resulting bam file was randomly down-sampled to a million reads, which were aligned against Zv9/danRer7 and read densities across genomic features were estimated for RNA-Seq-specific quality control metrics. Read mapping and quantification was performed using Salmon v. 1.2.1 [73], with the following flags: quant -l IU --validateMappings in paired-end mode against the GRCz11/danRer11 genome assembly and ENSEMBL 100 annotation, using all GRCz11 sequences as decoys. Quantification files were processed in R using the txImport function and per-gene counts and TPM estimates were retrieved [74].

#### *rbpms2*-null RNA-Seq strategy and analysis

RNA integrity and concentration were checked on a Fragment Analyzer (Advanced Analytical) and libraries were prepared using the Illumina NeoPrep RNAseq kit. Blue Pippin size selection was performed, selecting for 350 nt fragments. Libraries were quantified using the Fragment Analyzer (Advanced Analytical) and qPCR before being loaded for paired-end sequencing 2x75 nucleotides using the Illumina NextSeq 500 in High Output mode. For quality control purposes, reads were aligned against GRCz10/danRer10 (Sept. 2014) using bwa mem v. 0.7.12-r1039 with flags –t 16 –f and mapping rates, fraction of multiply-mapping reads, number of unique 20-mers at the 5' end of the reads, insert size distributions and fraction of ribosomal RNAs were calculated using dedicated perl scripts and bedtools v. 2.25.0.64 [72]. In addition, each resulting bam file was randomly down-sampled to a million reads, which were aligned against GRCz10/danRer10 and read density across genomic features were estimated for RNA-Seq-specific quality control metrics. RNA-Seq mapping and quantitation was done against GRCz10 / ENSEMBL 89 annotation using STAR v. 2.5.3a [75] with flags -runThreadN 8 --runMode alignReads --outFilterType BySJout --outFilterMultimapNmax 20 --alignSJoverhangMin 8 --alignSJDBoverhangMin 1 --outFilterMismatchNmax 999 --alignIntronMin 10 --alignIntronMax 1000000 --alignMatesGapMax 1000000 --outSAMtype BAM SortedByCoordinate --quantMode TranscriptomeSAM with --genomeDir pointing to a 75nt-junction GRCz10/danRer10 STAR suffix array. Gene expression was quantitated using RSEM v. 1.3.0 [76] with the following flags for all libraries: rsem-calculate-expression --calc-pme --alignments -p 8 --forward-prob 0 against an annotation matching the STAR SA reference. Posterior mean estimates (pme) of counts and estimated RPKM were retrieved. For both RNA-Seq experiments, differential expression analysis was performed using DESeq2 on count data comparing Nkx2.5+ vs. bulk and *rbpms2*-null vs. control samples, respectively. For *rbpms2*-null analyses, a batch correction was introduced in the form of a count ~ genotype+batch additive model, given substantial batch-related sample dispersion upon visual inspection of the first and second principal components of variance. No Cook's cutoff, independent filtering nor log-fold-change shrinkage were applied. Log2-fold changes as well as raw and Benjamini-Hochberg adjusted p-values were calculated for each protein-coding gene [77].

**Splicing analysis**

Analysis of differential RNA processing in *rbpms2*-null vs. control samples was performed in three distinct ways. Initially, isoform-level expression data were retrieved from RSEM and analyzed in DESeq2, using the posterior mean estimates of the counts per isoform as input and a similar batch-correction strategy as the gene-level differential-expression analysis described above. Additionally, the dedicated splicing analysis algorithm rMATS.3.2.5 was run on the data as RNASeq-MATS.py with flags - -t paired -len 75 -c 0.1 -analysis U -libType fr-firststrand -novelSS 1 -keepTemp pointing to the same GTF file and STAR index used at the read mapping star, across all samples and batches. Event inclusion was quantified both by counting junction reads only and junction and read on targets. The resulting data collated across replicates were filtered, and events had to meet the following conditions to be retained: the event had to be called in all samples, differences in inclusion levels had to be > 0.1 or < -0.1 and false-discovery rate < 0.1. Finally, the MISO algorithm was used to assess expression and usage of complex alternative isoforms of the pln gene and visualized with Sashimi plots [57,78]. Briefly, dedicated annotations of the isoforms of interest were generated manually and run using MISO v. 0.5.3 in the python 2.7 environment with default parameters and --read-len 75. MISO outputs were compared between genotypes within each batch using the compare_miso –compare-samples function.

**Whole mount in situ hybridization**

In situ hybridization was performed essentially as described [79]. Templates for generating anti-sense riboprobes for *rbpms2a, rbpms2b, mybpc3* isoform-specific*, mybpc3* total transcript*, pln* isoform-specific*,* and *pln* total transcript targets were generated by PCR or by gBlock (IDT) using the respective primer pairs or gBlock sequences shown in Table S6. All sequences were cloned into pCR4 vector using the TOPO TA cloning kit (Thermo Fisher Scientific). The *rbpms2a* and *rbpms2b* templates were linearized with BamHI and KpnI respectively, and Digoxygenin-labeled antisense probes were generated using T7 polymerase. Templates for *pln* isoform-specific, *pln* total transcript, and *mybpc3* total transcript probes were linearized with SpeI and transcribed using T7. The template for the *mybpc3* isoform-specific probe was linearized with NotI and transcribed with T3 polymerase. All probes were synthesized using the DIG RNA Labeling Kit (SP6/T7/T3; Millipore Sigma) and colorimetric reactions were performed using the NBT/BCIP chromogenic substrate (Promega Corp.).

**Whole mount immunofluorescence and confocal imaging**

Embryos were treated with 0.5M KCl to arrest hearts in diastole prior to fixation overnight in phosphate-buffered saline (PBS) containing 4% paraformaldehyde (PFA). The following day, embryos were thoroughly washed with PBS and dehydrated through a methanol series for storage at -20°C. Prior to staining, embryos were rehydrated into PBS containing 0.1% Tween-20 (PBST) and treated with bleaching solution (0.8% KOH, 0.9% $H_20_2$, 0.1% Tween-20) to remove pigment. Embryos were washed with PBST and permeabilized for 1 hour in PBS containing 0.5% Triton X-100. Prior to antibody incubation, embryos were blocked in 5% bovine serum albumin (BSA) and 5% goat serum in PBST for 1 hour. Embryos were incubated with primary antibodies diluted in blocking solution overnight at 4°C. The following primary antibodies were utilized: anti-Troponin T (CT3; Developmental Studies Hybridoma Bank (DSHB); 1:500 dilution), anti-GFP (B-2; Santa Cruz Biotechnology; 1:100), anti-RBPMS2 (ab181098; abcam; 1:500), anti-Tropomyosin (CH1; DSHB; 1:200) and anti-Myosin heavy chain (MF20; DSHB; 1:50). Embryos were washed in PBST and incubated with Alexa Fluor-conjugated secondary antibodies listed in the STAR methods table (1:500, Thermo Fisher Scientific) for 1-3 hours at room temperature. Following incubation, embryos were washed in PBST and nuclei labeled with DAPI for 5’ (Sigma-Aldrich; 1:5000). Embryo heads were then removed to expose the heart. Embryos were mounted in 0.9% low-melt agarose on glass-bottom dishes (MatTek Corp.) for imaging on a Nikon Ti Eclipse confocal microscope. Images were processed and/or analyzed for nuclei counts with ImageJ [80] Further processing was performed in Photoshop (Adobe Inc.).

**Western Blotting**

Western blotting was performed as previously described [81]. Lysates were prepared from pools of 5 embryos at 48 hpf and were probed with anti-RBPMS2 (ab181098; abcam; 1:1000) and anti-Alpha Tubulin (DM1A; Millipore; 1:10,000) overnight at 4°C. HRP-conjugated anti-rabbit (7074; Cell Signaling) and anti-mouse (7076; Cell Signaling) secondary antibodies were used at a 1:10,000 dilution.

**Histological sectioning and staining**

Paraffin sections of adult zebrafish hearts were generated following conventional procedures used for dissected mouse hearts. Briefly, hearts were dissected and fixed overnight in 4% PFA followed by washes in PBS and serial dehydration into 100% ethanol. Hearts were stored in 100% ethanol overnight at 4°C to and cleared with xylenes prior to paraffin embedding. 7μm sections were then transferred to microscope slides for histological or antibody staining. Prior to staining, slides were taken back through xylenes, 100% ethanol, and into water or PBS. Hematoxylin and Eosin (H&E) staining of paraffin sections were performed using conventional methodology and reagents (Sigma-Aldrich). Acid fuchsin/Orange G (AFOG) staining was performed as previously described [82] using Bouins solution (Thermo Fisher Scientific), aniline blue (Sigma-Aldrich), orange G (Sigma-Aldrich), and acid fuchsin (Sigma-Aldrich). H&E or AFOG stained sections were dehydrated in ethanol, washed with xylenes, and mounted with Cytoseal (Thermo Fisher Scientific) prior to imaging.  Immunostaining of paraffin sections was performed as previously described [83] using anti-RBPMS2 (ab181098; abcam; 1:500) and anti-Myosin heavy chain (MF20; DSHB; 1:50) antibodies followed by incubation with DAPI for 5'

(Sigma-Aldrich; 1:5000). Stained sections were then mounted with Fluorsave (Millipore) and kept in the dark.

**Assessments of cardiac function in zebrafish**

Volumetric measurements of the zebrafish heart were performed at ~48 hpf using light sheet fluorescence microscopy and the deep-learning neural network CFIN as described previously [42]. Briefly, live-mounted *Tg(myl7:GFP)* zebrafish were imaged with a 60X objective on a custom light sheet fluorescence microscope that was built based on a previously published design [84,85]. Hearts were imaged through the entire ventricle by collecting dynamic plane images at each GPF-positive z depth. Dynamic images were taken every 1µm and spanned at least four cardiac cycles. In order to ensure optimal CFIN performance with images acquired on a custom-built microscope, a subset of these images were manually labeled and used to retrain the network as previously described. Experimental images were then processed and analyzed by CFIN in MATLAB (MathWorks). Fractional shortening (FS) in 33 hpf hearts was determined from videos of *Tg(myl7:GFP)* zebrafish hearts. One-dimensional measurements of the arterial pole width during diastole (d) and systole (s) were obtained in ImageJ and used to calculate percent FS.

**Reverse transcription quantitative polymerase chain reaction (RT-qPCR)**

Total RNA was harvested from pooled material (10 embryos or 10 dissected hearts per pool) using TRIzol reagent and purified via Direct-zol RNA MicroPrep columns (ZYMO). cDNA synthesis was performed using Superscript III (Invitrogen). RT-qPCR was performed on the QuantStudio 3 Real-Time PCR system (Applied Biosystems) using KAPA SYBR FAST qPCR master mix reagents (Kapa Biosystems). Relative mRNA expression was determined after normalizing to *rps11* using the $\Delta\Delta$Ct method [86]. 3 biological replicates (3 separate pools of 10 embryos each) and 3 technical replicates were performed for each experiment. Statistical significance was determined by an unpaired, two-tailed Student's t-test assuming equal variances.

**Calcium imaging of zebrafish hearts**

Hearts were isolated manually from 48 hpf zebrafish embryos and placed in normal Tyrode's (NT) solution [NT contains 136mM NaCl, 5.4mM KCl, 1mM $MgCl_2x6H_2O$, 5mM D-(+)-Glucose, 10mM HEPES, 0.3mM $Na_2HPO_4x2\ H_2O$, 1.0mM $CaCl_2x2\ H_2O$, pH 7.4] supplemented with 8 mg/ml BSA. For ratiometric Ca2+-transient recordings, hearts were loaded for 15 minutes with 50µM of the Ca2+-sensitive dye Fura-2, AM (Thermo Fisher Scientific) and subsequently incubated in due-free NT buffer with BSA for 45' to allow complete intracellular hydrolysis of the esterified dye. Individual hearts were loaded into a perfusion chamber (RC-49MFS, Warner Instruments) mounted on an inverted microscope (TW-2000, Nikon). The perfusion chamber contained NT supplemented with 1mM Cytochalasin D to inhibit cardiac contraction. A high-speed monochromator (Optoscan, Cairn Research) was used to rapidly switch the excitation wavelength between 340nm and 380nm with a bandwidth of 20nm and at a frequency of $500s^{-1}$. The excitation light, generated by a 120W metal halide lamp (Exfo X-Cite 120, Excelitas Technologies) was reflected by a 400nm cutoff dichroic mirror and fluorescence emission was collected by the camera through a 510/580nm emission filter. For the measurement of fluorescence intensities, we used a high-speed 80×80-pixel CCD camera (CardioCCD-SMQ, RedShirtImaging, LLC) with 14-bit resolution. For ratiometric Ca2+-transient recordings, monochromator and camera were synchronized. Because each wavelength change required a mechanical movement of the grating, each ratio required the recording of four frames, where one frame was used for each transition between wavelengths. This ultimately resulted in the acquisition of ratiomateric Ca2+ measurements at a frequency of 125 Hz. Measurements were performed using a 40x objective and a 0.28x C-mount adapter (final magnification was 11.2x), resulting in a pixel-to-pixel distance of 2.14μm. Images were analyzed and Ca2+ transient durations, amplitudes and diastolic Ca2+ were quantified with MATLAB (Mathworks) using customized software as previously described [87]. The values plotted in Fig. 6 represent recordings from the center of each chamber.

**hiPSC-CM Differentiation**

hiPSC were seeded onto 12-well plates at 60,000 cells/well and maintained for 48 hours prior to differentiation or until they reached 60% confluence. On days 0 – 2 of differentiation, cells were cultured with RPMI (Cat. # 11875093, Thermo Fisher Scientific) supplemented with B27 minus insulin (Cat. # A1895601, Thermo Fisher Scientific) and 6μM CHIR99021 (Cat. # 72054, Stem Cell Technologies). Cells were replenished with RPMI/B27(-insulin) on day 3, then cultured with RPMI/B27(-insulin) supplemented with 5μM IWR1-endo (Cat. # 2564, Stem Cell Technologies) on days 4 and 5. From days 6 – 12, cells were maintained in RPMI/B27(-insulin) with media changes every 48 hours. Lactate selection was performed on day 12 for 48hrs with RPMI supplemented with 5mM sodium lactate (Cat. # L7022, MilliporeSigma). For downstream experiments, iPSC-CMs were dissociated using StemPro Accutase (Cat. A1110501, Thermo Fisher Scientific). Cells were gently resuspended with an equal volume of RPMI/B27 supplemented with 5% FBS and 5μm ROCK inhibitor (Y-27632, Tocris), passed through a 70μm cell strainer and pelleted at 200 x g for 5 min.

**Immunostaining for hiPSC-CMs**

hiPSC-CMs were dissociated as stated above and re-plated on glass cover slips coated in GelTrex. Cells were allowed to recover in RPMI/B27 with 5% FBS and 5μm ROCK inhibitor overnight followed by replacement with RPMI/B27 every 48 hours until fixation. Cells were fixed in 4% PFA (10 minutes) and permeabilized in PBS with 0.1% Triton X-100 (5 minutes). Cells were blocked in PBS with 3% BSA and incubated with primary antibodies overnight at 4°C. The following primary antibodies were utilized: anti-Sarcomeric alpha actinin (ab9465; abcam; 1:500) and anti-Troponin T (CT3; DSHB; 1:500) followed by DAPI staining to mark nuclei. Stained coverslips were mounted with FluorSave (Millipore) and stored in the dark.

**Calcium imaging of hiPSC-CMs**

Single line scanning confocal microscopy was used to record calcium transients and calcium release events. In brief, 100,000 hiPSC-CMs were seeded onto 8mm GelTrex-coated coverslips and cultured for 4 days. Cells were loaded with 1uM fluo-4 calcium indicator (Cat. F14201; Thermo Fisher Scientific) prepared in Tyrode's solution [137mM NaCl, 2.7mM KCl, 1mM $MgCl_2$, 1.8mM $CaCl_2$, 0.2mm $Na_2$-$HPO_4$, 12mM $NaHCO_3$, 5.5mM D-Glucose, pH7.4] for 10 minutes at 37°C. Cells were then washed with DPBS and incubated in Tyrode's solution for 10 minutes prior to imaging. Temperature-controlled calcium measurements were acquired for a total duration of 30s using the following imaging protocol: 10s spontaneous recording, 10s paced, 10s recovery. Cells were electrically paced at 1hz with 15V using a MyoPacer (IonOptix). Multiple line scans were compiled in Image J [80] using a custom macro. Calcium transient kinetics were analyzed using MATLAB (MathWorks).

**SUPLEMENTAL FIGURES**

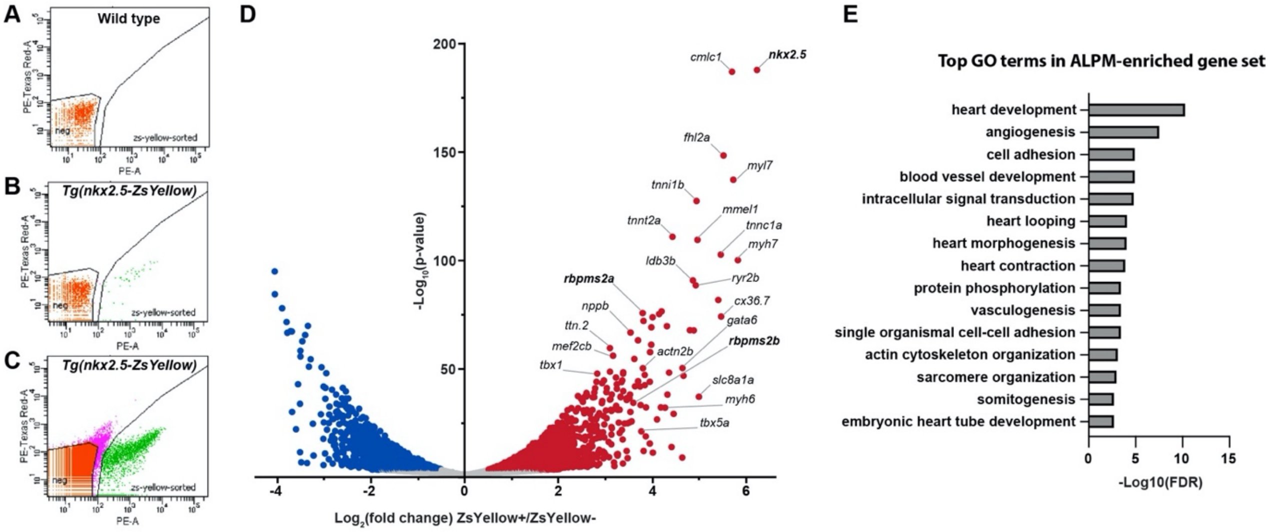


**Figure S1 – Fluorescence activated cell sorting, RNA-sequencing, and differential expression analysis of ZsYellow-positive cardiopharyngeal progenitors and early-differentiating cardiomyocytes in the anterior lateral plate mesoderm of *Tg(nkx2.5:ZsYellow)* embryos.** (A-C) Flow-cytometry dot plots showing the fluorescence intensities of single cells dissociated from 14-16 somite stage (ss) wild-type control (CTRL; A) and *Tg(nkx2.5:ZsYellow)* (B,C) embryos in the yellow (PE-A; x-axis) and red (PE-Texas Red-A; y-axis) channels. The experiment shown in (A,B) was used to set the gates for purifying both ZsYellow-positive (zs-yellow-sorted) and ZsYellow-negative (neg) cells by fluorescence activated cell sorting [representative sort plot shown in (C)]. (D) Volcano plot showing the distribution of fold change (FC) and raw p-values for protein-coding RNAs expressed in ZsYellow-positive cells relative to ZsYellow-negative cells. Transcripts meeting the inclusion criteria for Gene Ontology (GO) term analysis ($|FC|>1.5$; adjusted $p\text{-value}<0.05$) are highlighted in purple ($FC<-1.5$) or red ($FC>1.5$). Transcription factors and myocardial genes known to mark the lateral plate mesoderm in zebrafish or higher vertebrates are labelled. *nkx2.5*, *rbpms2a*, and *rbpms2b* are shown in bold because we used nkx2.5 promoter activity to distinguish the two populations, and *rbpms2a* and *rbpms2b.* (E) Bar graph showing the top GO terms (BP-direct; $FC>1.5$; adjusted $p\text{-value}<0.05$) ranked by decreasing false discovery rate (FDR). The complete list of GO terms is reported in Table S1.

**A**

```
                                                                   RRM
Human RBPMS2        MSNLKPDGEHGGSTGTGSGAGSGGALEEEVRTLFVSGLPVDIKPRELYLLFRPFKGYEGS 60
Zebrafish Rbpms2a   -----------MSLKSDSETNTSVSLEEEVRTLFVSGLPVDIKPRELYLLFRPFKGYEGS 49
Zebrafish Rbpms2b   -----------MSVKSDSEPNNNVSLEEEVRTLFVSGLPTDIKPRELYLLFRPFKGYEGS 49
                                *  :.*  ... :**************.********************

                                   RRM
Human RBPMS2        LIKLTARQPVGFVIFDSRAGAEAAKNALNGIRFDPENPQTLRLEFAKANTKMAKSKLMAT 120
Zebrafish Rbpms2a   LIKLTSKQPVGFVTFDSRSGAEAAKNALNGIRFDPESPQTLRLEFAKANTKMAKSKLMAT 109
Zebrafish Rbpms2b   LIKLTSKQPVGFVTFDSRSGAEAAKNALNGVRFDPENPQTLRLEFAKANTKMAKSKLMGT 109
                    *****::****** ****:***********:*****.*********************.*

Human RBPMS2        PNPSNVHPALGAHFIARDPYDLMGAALIPASPEAWAPYPLYTTELTPAISHAAFTYPTAT 180
Zebrafish Rbpms2a   PNPSNLHPALGAHFIARDPYDLTGAALIPASPEAWAPYPLYTTELTPGLPHAAFTYPAAA 169
Zebrafish Rbpms2b   PNPTNIHPALGAHFIARDPYDLTGAALIPASPEAWSPYPLYTPELSPGLPHTAFTYPAAA 169
                    ***:*:**************** ************:****** **:*.: *:*****:*:

Human RBPMS2        A-AAAALHAQVRWYPSS-DTTQQGWKYRQFC 209
Zebrafish Rbpms2a   A-AAAALHAQMRWYPSPSESSQPGWKSRQFC 199
Zebrafish Rbpms2b   AAAAAALHAQMRWYPSTSDSSQPGWKSRQFC 200
                    * ********:*****  :::* *** ****
```

**B**

| Human vs. | Full Length | RRM |
|---|---|---|
| Rbpms2a | 84.3% | 93.6% |
| Rbpms2b | 81.3% | 92.3% |

**Figure S2 – Amino acid sequence conservation between zebrafish and human RBPMS2 proteins.** (A) Alignment of human and zebrafish RBPMS2 proteins created with Clustal Omega. Asterisks indicate conserved amino acids, colons indicate partially conserved or strong similarities in amino acid properties, and periods indicate amino acids with weak similarity. RNA-recognition motifs (RRM), as predicted by the Prosite and InterPro databases, are highlighted in grey. (B) Table showing the percent identities between zebrafish Rbpms2a or Rbpms2b and human RBPMS2 across either the full-length sequences or within the RRMs.

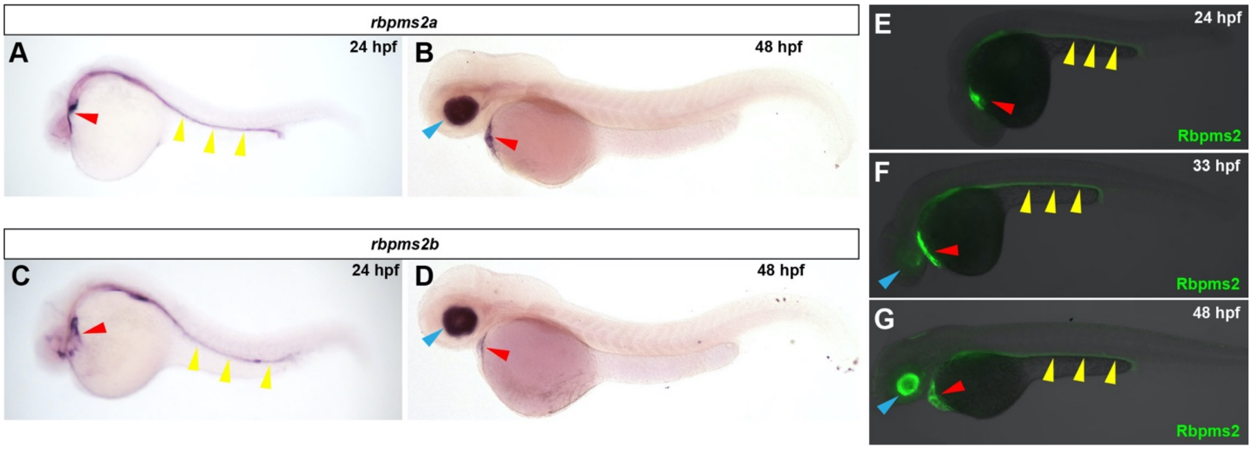


**Figure S3 - Developmental expression patterns of Rbpms2a and Rbpms2b in zebrafish.** (A-D) Brightfield images of 24 hours post fertilization (hpf; A,C), and 48 hpf (B-D) embryos processed for in-situ hybridization with *rbpms2a* (A-B) or *rbpms2b* riboprobes (C-D). (E-G) Merged brightfield and fluorescent images of 24 hpf (E), 33 hpf (F), and 48 hpf (G) embryos immunostained with an antibody that detects Rbpms2. Red, yellow, and blue arrowheads highlight staining in the heart, pronephric duct, and retina, respectively. For all experiments, little to no variation in expression patterns were observed between animals in each group (n>10/group).

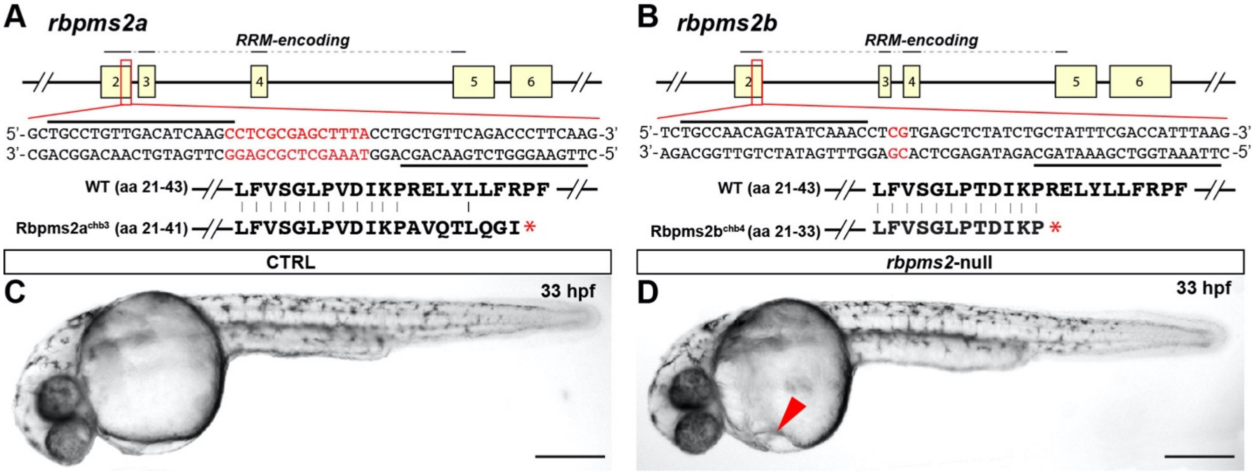


**Figure S4 - Properties of the *rbpms2a*$^{chb3}$ and *rbpms2b*$^{chb4}$ mutant alleles.** (A,B) Schematic diagrams (top) of the zebrafish *rbpms2a* and *rbpms2b* loci (partial; exons 2-6; top) showing the exon regions encoding the RNA-recognition motifs (RRMs; solid lines). DNA sequences (middle) surrounding and including the TALEN target sites (black lines) in exon 2 of both loci and nucleotides deleted in the *rbpms2a*$^{chb3}$ (A) and *rbpms2b*$^{ch4}$ alleles (B; red). Partial amino acid sequences (bottom) of Rbpms2a and Rbpms2b aligned with the corresponding sequences of the predicted protein products of *rbpms2a*$^{chb3}$ and *rbpms2b*$^{chb4}$. Vertical lines highlight identical amino acids. The asterisk shows the location of a premature stop codon in the shifted reading frame. (C,D) Brightfield images of 33 hours post fertilization control-sibling (CTRL; C) and *rbpms2*-null (D) zebrafish embryos. Red arrowhead highlights pericardial edema in *rbpms2*-null animals. Little to no variation in phenotype was observed between animals in each group (n>10/group). Scale bars=250 μm.

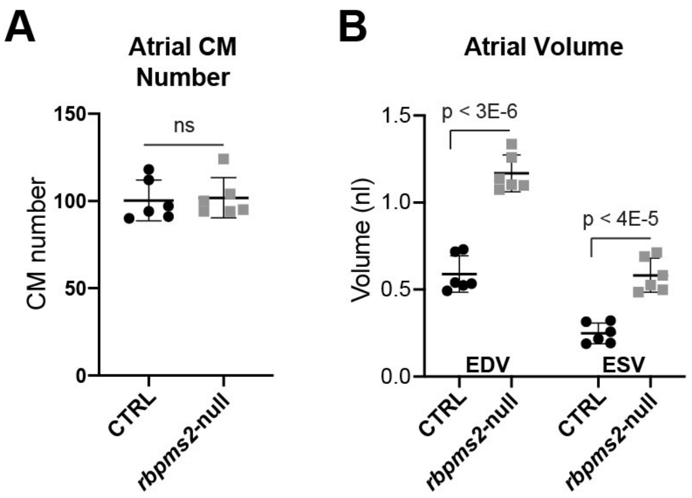


**Figure S5 - Atrial cardiomyocyte number and volume in *rbpms2*-null animals.** (A) Dot plot showing atrial cardiomyocyte number in control-sibling (CTRL; n=6) and *rbpms2*-null (n=6) animals at 48 hours post fertilization (hpf). (B) Dot plots showing the end-diastolic volume (EDVs) and end-systolic volume (ESVs) for the atrium of 48 hpf CTRL (n=6) and *rbpms2*-null (n=6) animals as determined by the Cardiac Functional Imaging Network (CFIN). Error bars show one standard deviation. Statistical significance was determined by an unpaired, two-tailed Student's t-test assuming equal variances. ns, not significant.

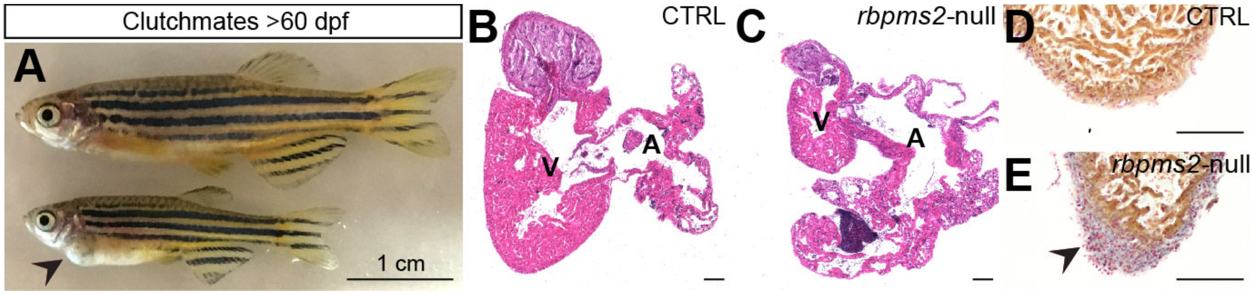


**Figure S6 - Cardiac phenotype of late-juvenile or adult zebrafish.** (A) Brightfield image of control-sibling (top) and *rbpms2*-null zebrafish (bottom) from the same clutch after >60 days post fertilization (dpf). Little to no variation was observed in the gross phenotypes of *rbpms2*-null animals living beyond 60 dpf (n>10). Black arrowhead highlights cardiac edema in the double mutant. (B-E) Brightfield images of cardiac sections from control-sibling (CTRL; B,D) and *rbpms2*-null (C,E) animals stained with Hematoxylin and Eosin (B,C) or Acid Fuchsin Orange G (D,E). Little to no variation was observed between 27 sections examined per histological stain from each experimental group (3 hearts/group; 9 sections/heart). Black arrowhead highlights myocardial fibrosis in the double-mutant heart. Scale bars, 1 cm, (A); 100 μm, (B-E).

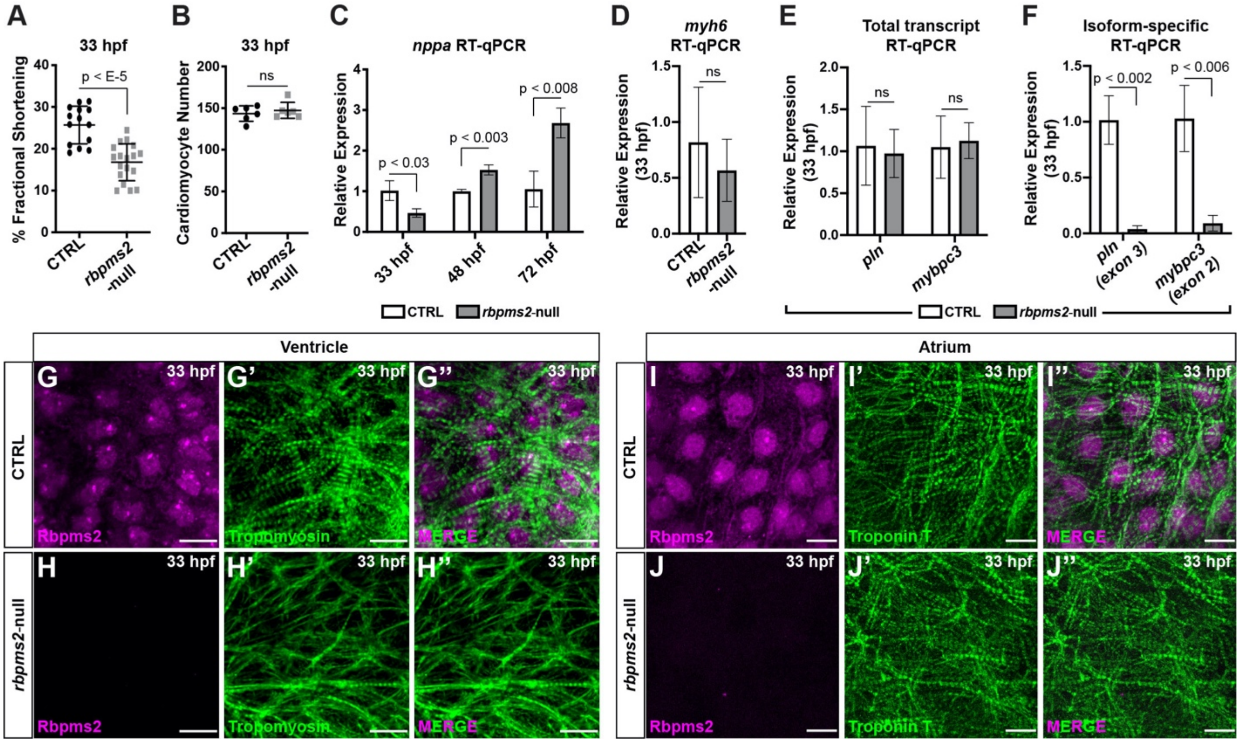


**Figure S7 – Early-stage phenotypic analysis of *rbpms2*-null animals.** (A) Dot plot showing the percent fractional shortening at the arterial pole of the heart tube in control (CTRL) and *rbpms2*-null embryos at 33 hours post fertilization (hpf). (B) Dot plot showing total cardiomyocyte numbers in CTRL (n=6) and *rbpms2*-null (n=6) embryos at 33 hpf. (C) Bar graph showing the relative expression levels of *nppa* in CTRL and *rbpms2*-null embryos at 33 hpf, 48 hpf, and in dissected 72 hpf hearts. (D) Bar graph showing the relative expression levels of *myh6* in CTRL and *rbpms2*-null embryos at 33 hpf. (E,F) Bar graphs showing the relative expression levels of the total (E) and isoform-specific (F) *mybpc3* and *pln* populations in CTRL and *rbpms2*-null embryos at 33 hpf. For (A-F), error bars show one standard deviation, and statistical significance was determined by an unpaired, two-tailed Student's t-test assuming equal variances. ns, not significant. Scale bars=10 µm.

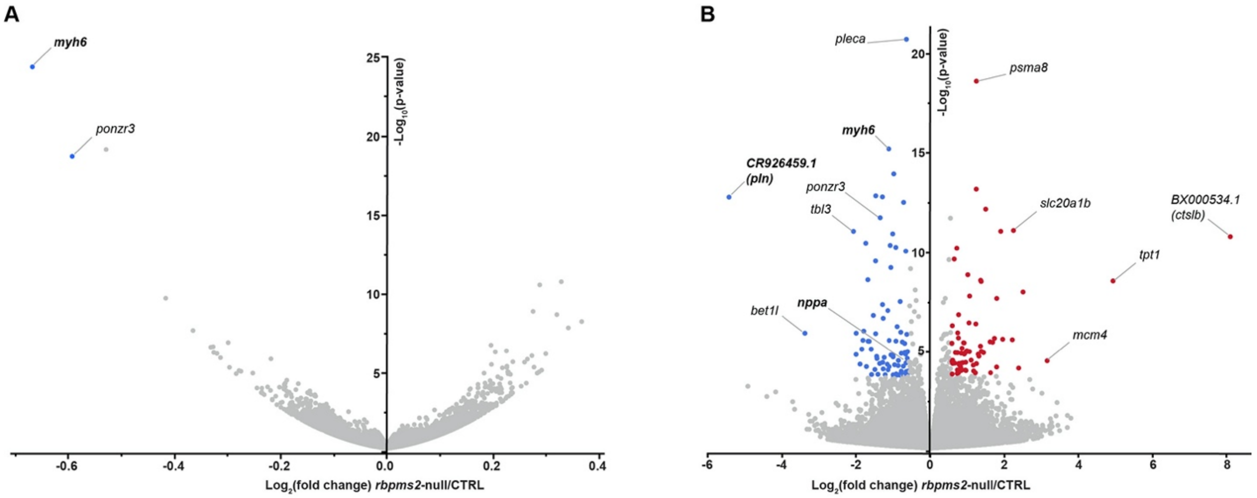


**Figure S8 - Gene-level and isoform-level differential expression analysis of control-sibling and *rbpms2*-null animals at 33 hours post fertilization.** (A,B) Volcano plots showing the distribution of fold change and raw p-values for genes (A) or gene isoforms (B) expressed in 33 hours post fertilization *rbpms2*-null animals compared to control-sibling embryos. Transcripts meeting the inclusion criteria for Gene Ontology (GO) term analysis (|FC|>1.5; adjusted p-value<0.05) are highlighted in purple (FC<-1.5) or red (FC>1.5). The gene names shown in bold font were validated by qPCR or in situ hybridization.

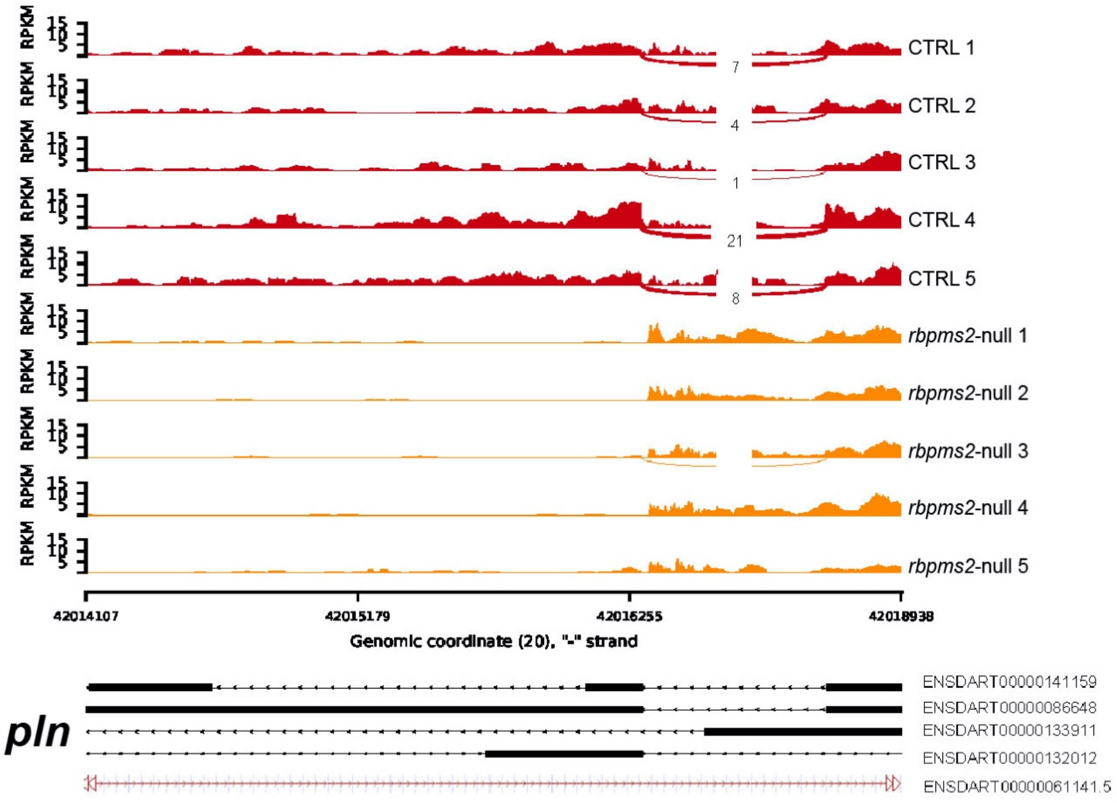


**Figure S9 – Mixture of Isoforms (MISO) splicing analysis across the *pln* locus.** Alternative 3' exon usage in the *pln* locus in sibling control (CTRL) and *rbpms2*-null animals. RNA-sequencing read densities and junction read configurations across the 3' end of the gene were quantified and plotted using MISO and Sashimi, and the results are compatible with genotype-specific regulation of isoform use (danRer10 genomic coordinates, ENSEMBL 91 annotation).

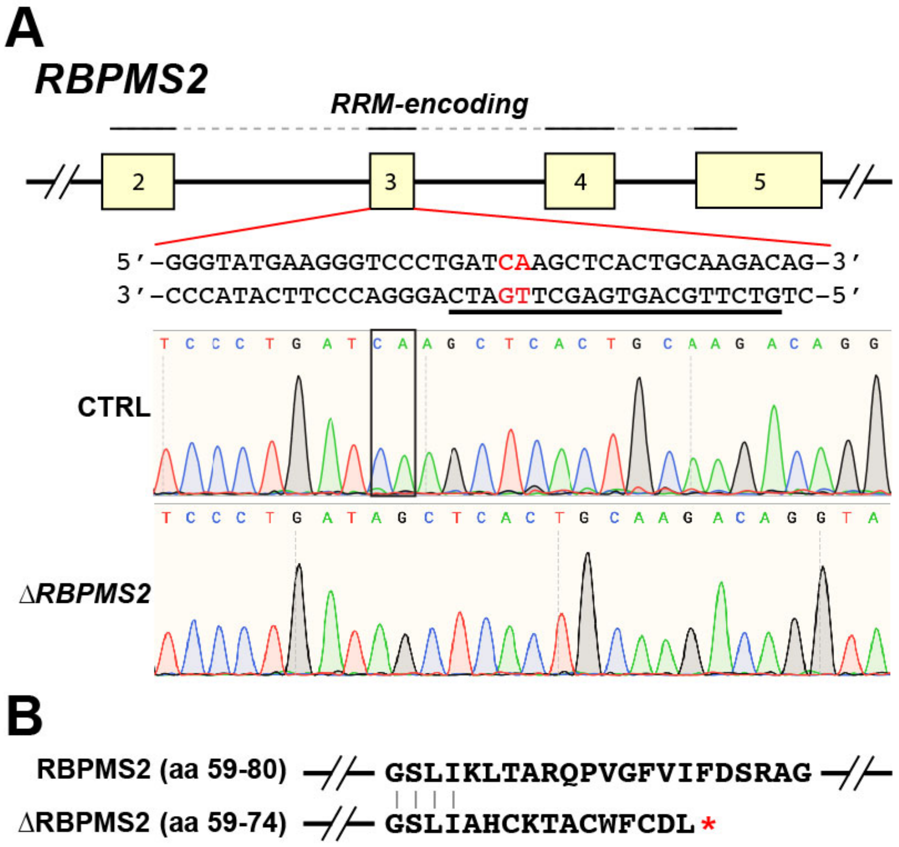


**Figure S10 - Properties of the *ΔRBPMS2* null allele created in human induced pluripotent stem cells**. (A) Schematic diagram (top) of the human *RBPMS2* locus (partial; exons 2-5) highlighting the exon regions encoding the RNA recognition motif (RRM; solid black lines). DNA sequence (middle) of *RBPMS2* exon 3 highlighting the guide RNA sequence (black line) and two nucleotides deleted in *ΔRBPMS2* (red). Sanger sequencing chromatograms (bottom) showing *RBPMS2* exon 3 sequences in the parental (CTRL) and *ΔRBPMS2* strains of induced pluripotent stem cells. The black box highlights the two nucleotides in the parental strain, which are deleted in both *RBPMS2* alleles of the *ΔRBPMS2* strain. (B) Partial amino acid sequence of wild-type RBPMS2 (top) and aligned sequence of the ΔRBPMS2 predicted protein product (bottom). Vertical lines highlight identical amino acids. The asterisk shows the location of a premature stop codon in the shifted reading frame.

**Table S1** – Differentially expressed genes and associated Gene Ontology terms in ZsYellow-positive cells relative to ZsYellow-negative cells purified from 14-16 somite stage *Tg(nkx2.5:ZsYellow)* zebrafish embryos.

**Table S2** – Differentially expressed genes and associated Gene Ontology terms in *rbpms2*-null embryos relative to control siblings at 33 hours post fertilization.

**Table S3** – Differentially expressed gene isoforms and associated Gene Ontology terms in *rbpms2*-null embryos relative to control siblings at 33 hours post fertilization.

**Table S4** – Differentially alternatively-spliced genes in *rbpms2*-null embryos relative to control siblings at 33 hours post fertilization identified by rMATS.

**Table S5** – MISO analysis for alternative splicing of *pln* in *rbpms2*-null embryos relative to control siblings at 33 hours post fertilization.

**Table S6** – Primer and other oligonucleotide sequences.
